# PTEN regulates adipocyte progenitor growth, differentiation and replicative aging

**DOI:** 10.1101/2021.01.05.425433

**Authors:** Anna S. Kirstein, Stephanie Kehr, Michèle Nebe, Martha Hanschkow, Judith Lorenz, Melanie Penke, Jana Breitfeld, Diana Le Duc, Kathrin Landgraf, Antje Körner, Peter Kovacs, Peter F. Stadler, Wieland Kiess, Antje Garten

## Abstract

The tumor suppressor phosphatase and tensin homolog (PTEN) negatively regulates the insulin signaling pathway. Germline PTEN pathogenic variants cause PTEN Hamartoma Tumor Syndrome (PHTS), associated with lipoma development in children. It remains unclear which mechanisms trigger this aberrant adipose tissue growth. Adipocyte progenitor cells (APCs) lose their capacity to differentiate into adipocytes during continuous culture, while APCs from PHTS patients’ lipomas retain their adipogenic potential over a prolonged period. To investigate the role of PTEN in adipose tissue development we performed functional assays and RNA sequencing of control and PTEN knockdown APCs. Reduction of PTEN levels using siRNA or CRISPR lead to an enhanced proliferation and differentiation of APCs. FOXO1 was downregulated on the mRNA level while inactivation through phosphorylation increased. FOXO1 phosphorylation initiates the expression of the lipogenesis activating transcription factor SREBP1. SREBP1 levels were higher after PTEN knockdown and may account for the enhanced adipogenesis. To validate this we overexpressed constitutively active FOXO1 in PTEN CRISPR cells and found reduced adipogenesis, accompanied by a SREBP1 downregulation. We observed that PTEN levels were upregulated during long term culture of wild type APCs. PTEN CRISPR cells showed less senescence compared to controls and the senescence marker CDKN1A (p21) was downregulated in PTEN knockdown cells. Cellular senescence was the most significantly enriched pathway found in RNA sequencing of PTEN knockdown vs. control cells. These results provide evidence that PTEN is involved in the regulation of APCs proliferation, differentiation and senescence, thereby contributing to aberrant adipose tissue growth in PHTS patients.

## Introduction

Adipose tissue distribution is altered during aging – while subcutaneous fat depots decrease in advanced age, fat accumulates in muscle, liver and bone marrow, leading to metabolic dysregulation (1). Responsiveness to insulin declines in adipocytes from older individuals and metabolic features related to fatty acid metabolism change (2, 3). Human adipose tissue contains mesenchymal stem cells (MSCs), which serve as adipocyte progenitors and contribute to adipose tissue regeneration throughout life (4). These cells are found within the stromal vascular fraction (SVF) of adipose tissue. Adipogenesis can be induced by insulin and other soluble factors *in vitro* (5). SVF cells from older individuals have a lower capacity for adipocyte differentiation (6) and during long term SVF cell culture the adipogenic potential declines (7). Inhibiting the phosphoinositide 3-kinase (PI3K)/AKT pathway in adipose progenitors using the mammalian target of rapamycin (mTOR) inhibitor rapamycin (8) or the PI3K inhibitor alpelisib (9) was shown to repress adipogenesis. Several studies link insulin signaling and aging. Mice with adipose tissue specific insulin receptor knockout had an increased lifespan (10), but the underlying mechanisms are controversial (11). Adipose tissue in these mice maintains mitochondrial activity and insulin-sensitivity during aging, indicating that insulin-sensitivity dynamics rather than insulin-resistance correlate with longevity (12, 11).

We observed that lipoma cells from a patient with a *phosphatase and tensin homolog* (*PTEN*) germline pathogenic variant retain their differentiation capacity over a prolonged period (13). PTEN is a lipid and protein phosphatase, mainly catalyzing the dephosphorylation of the second messenger phosphatidylinositol-3,4,5-trisphosphate, resulting in a deactivation of the PI3K/AKT pathway signaling. PTEN is an important antagonist of this pathway, which is activated by a multitude of extracellular signals including insulin and insulin-like growth factor 1 (IGF-1). PI3K/AKT signaling generally promotes cellular growth and survival. The downstream target and central signaling molecule mTOR is a major regulator of protein and lipid synthesis, cell growth, proliferation, autophagy and metabolism (14). Loss of the tumor suppressor *PTEN* is common in cancer. *PTEN* haploinsufficiency caused by germline pathogenic variants leads to the rare genetic disease PTEN hamartoma tumor syndrome (PHTS). PHTS patients show a wide variety of phenotypes including hamartomas of the skin, breast and thyroid, intestinal polyps, macrocephaly, vascular malformations and lipoma formation (15). Widespread abdominal lipomatosis and lack of subcutaneous adipose tissue was observed in a boy with PHTS (16). It remains unclear which specific factors cause this localized adipose tissue overgrowth in PHTS patients.

Several mouse models with *PTEN* downregulation in adipose tissue (17, 18), adipose progenitor subpopulations (17, 18) or osteoblast progenitors (19, 20) display adipose tissue redistribution and/or lipoma formation and partly recapitulate the human phenotype of PHTS. Overexpression of AKT in zebrafish also leads to lipoma formation, linking PI3K signaling to adipose tissue overgrowth (21). A high PTEN expression in adipose tissue (22) points to its importance in regulating normal adipose tissue function. *PTEN* pathogenic variants were found to lead to adipose tissue redistribution in mice (17, 18), with similar phenotypes also observed in humans (16).

To investigate the effects of PTEN downregulation in human adipose progenitor cells and create an *in vitro* model for PTEN insufficiency as seen in PHTS, we used SVF cells isolated from adipose tissue of healthy donors and downregulated PTEN via siRNA or clustered regularly interspaced short palindromic repeats (CRISPR) system. We thereby observed phenomena associated with proliferation, differentiation and replicative aging of fat cell progenitors pointing to a role for PTEN in lipoma formation.

## Results

### 2.1 PTEN downregulation enhanced PI3K signaling and SVF cell proliferation

To examine the impact of PTEN loss on adipocyte development we performed siRNA mediated knockdown of PTEN (PTEN KD) in SVF cells from visceral and subcutaneous adipose tissue of donors without *PTEN* mutation. As determined via Western blot analysis PTEN was reduced in the visceral siRNA knockdown cells to 0.49 ± 0.04 fold (p<0.0001, Figure 1a). On the mRNA level we found a reduction of *PTEN* expression to 0.33 ± 0.08 fold (p<0.0001) in visceral (Figure 1b) and 0.3 ± 0.01 fold (p=0.009) in subcutaneous SVF cells (Figure S1A). The knockdown was stable during 7 days of proliferation (Figure S1B) and 8 days of differentiation (Figure S1C). To test the functional significance of the knockdown we investigated activation of downstream PI3K/AKT pathway components via phosphorylation. Phosphorylated AKT (pAKT (T308)) was elevated 22 ± 14 fold (p=0.029) and ribosomal protein S6 phosphorylation (pS6 (Ser235/236)) was increased 13.0 ± 5.5 fold (p=0.0008) in visceral PTEN KD cells (Figure 1a).

**Figure 1:**
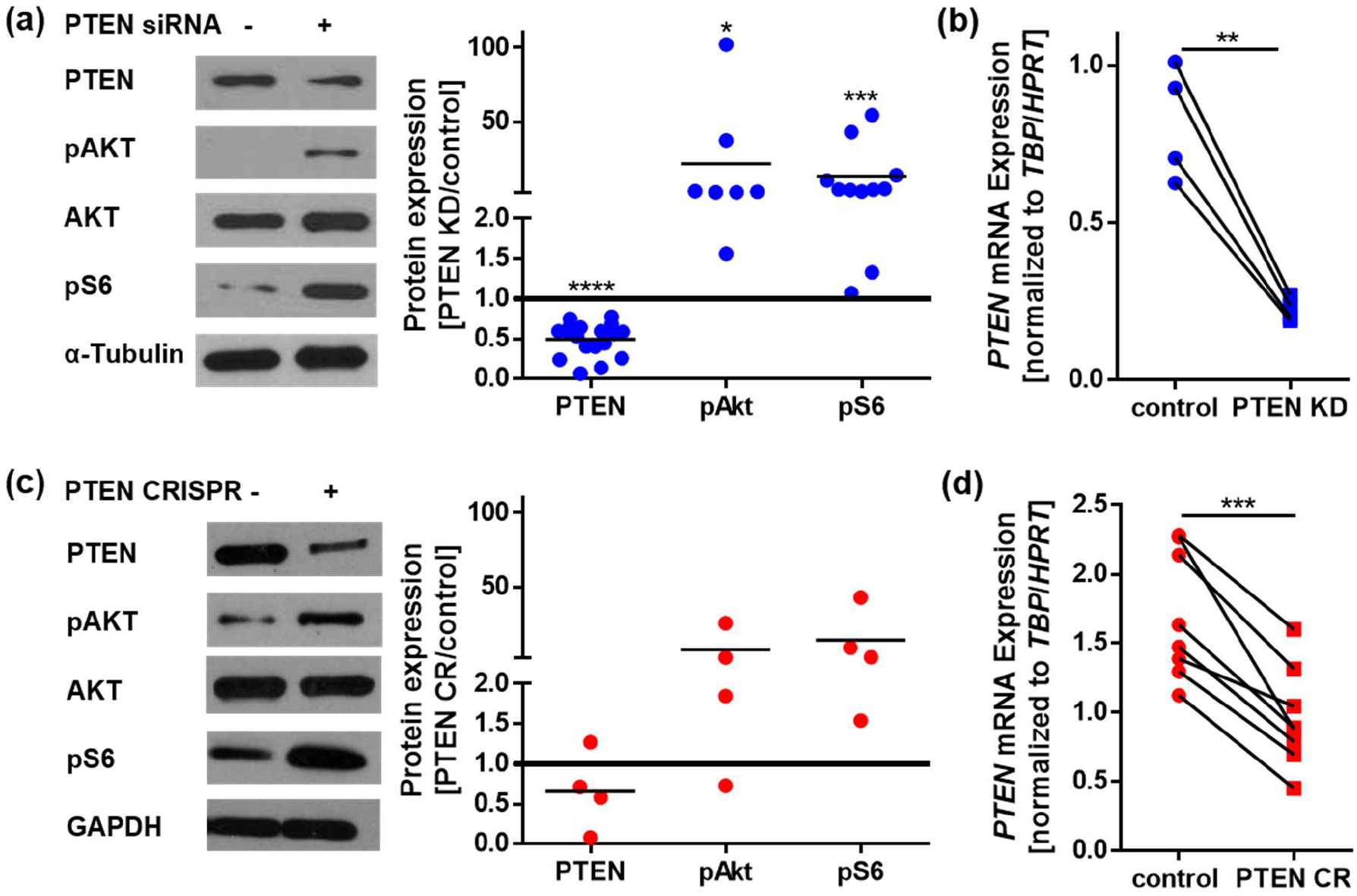
PTEN downregulation enhances PI3K signaling. (a) Western blots of control and PTEN siRNA (PTEN KD) transfected visceral SVF cells: PTEN was reduced to 0.49 ± 0.05 fold (normalized to α-Tubulin, n=18, p<0.0001), phosphorylated AKT (pAKT (T308)) was elevated 22 ± 14 fold (normalized to total AKT, n=7, p=0.029) and ribosomal protein S6 phosphorylation (pS6 (Ser235/236)) was increased 13.03 ± 5.5 fold (normalized to α-Tubulin, n=11, p=0.0008). (b) *PTEN* mRNA expression in control and PTEN siRNA transfected visceral SVF cells: *PTEN* was reduced to 0.33 ± 0.08 fold (normalized to means of *hypoxanthine phosphoribosyltransferase* (*HPRT*) and *TATA-box binding protein (TBP*) expression, n=4, p=0.0039). (c) Western blots of control and PTEN CRISPR (PTEN CR) SVF cells: PTEN was reduced to 0.7 ± 0.3 fold (normalized to α-Tubulin, n=4, p=0.27), pAKT (T308) was increased 8 ± 6 fold (normalized to total AKT, n=4, p=0.24) and pS6 (Ser235/236) was increased 14 ± 10 fold (normalized to α-Tubulin, n=4, p=0.09). (d) *PTEN* mRNA expression in control and PTEN CR cells: *PTEN* was reduced to 0.56 ± 0.05 fold (p=0.0002) (normalized to *HPRT* and *TBP*, n=4, p=0.0002). Lines between individual data points indicate matched data from single experiments (control vs. respective knockdown). p-values for (a) and (c) were determined via one-sample t-test of the log(fold change), p-values for (b) and (d) were determined via paired t-test.

siRNA mediated knockdowns are of transient nature, which is why we additionally performed *PTEN* knockouts using the CRISPR system in visceral SVF cells (PTEN CR). On average, PTEN protein levels were reduced to 0.7 ± 0.3 fold (p=0.27), while pAKT (T308) was increased 8 ± 6 fold (p=0.24) and pS6 (Ser235/236) was increased 14 ± 10 fold (p=0.09) (Figure 1c). *PTEN* mRNA was reduced to 0.56 ± 0.05 fold (p=0.0002) in PTEN CR cells (Figure 1d). Using both methods, we achieved a similar reduction in PTEN levels, as seen in cells of a patient with germline heterozygous *PTEN* deletion (13).

We observed an enhanced proliferation in PTEN insufficient SVF cells. PTEN downregulation led to faster expansion to a similar extent in PTEN KD cells (1.4 ± 0.2 fold, p=0.038 in visceral, 1.21 ± 0.08 fold, p=0.0006 in subcutaneous SVF cells, Figure 2a) and in PTEN CR cells (1.5 ± 0.1 fold, p=0.039, Figure 2b). The higher cell count was also reflected in a 1.09 ± 0.01 fold (p=0.0043) higher fraction of proliferation marker Ki-67 positive cells as shown by immunofluorescence staining (Figure 2c).

**Figure 2:**
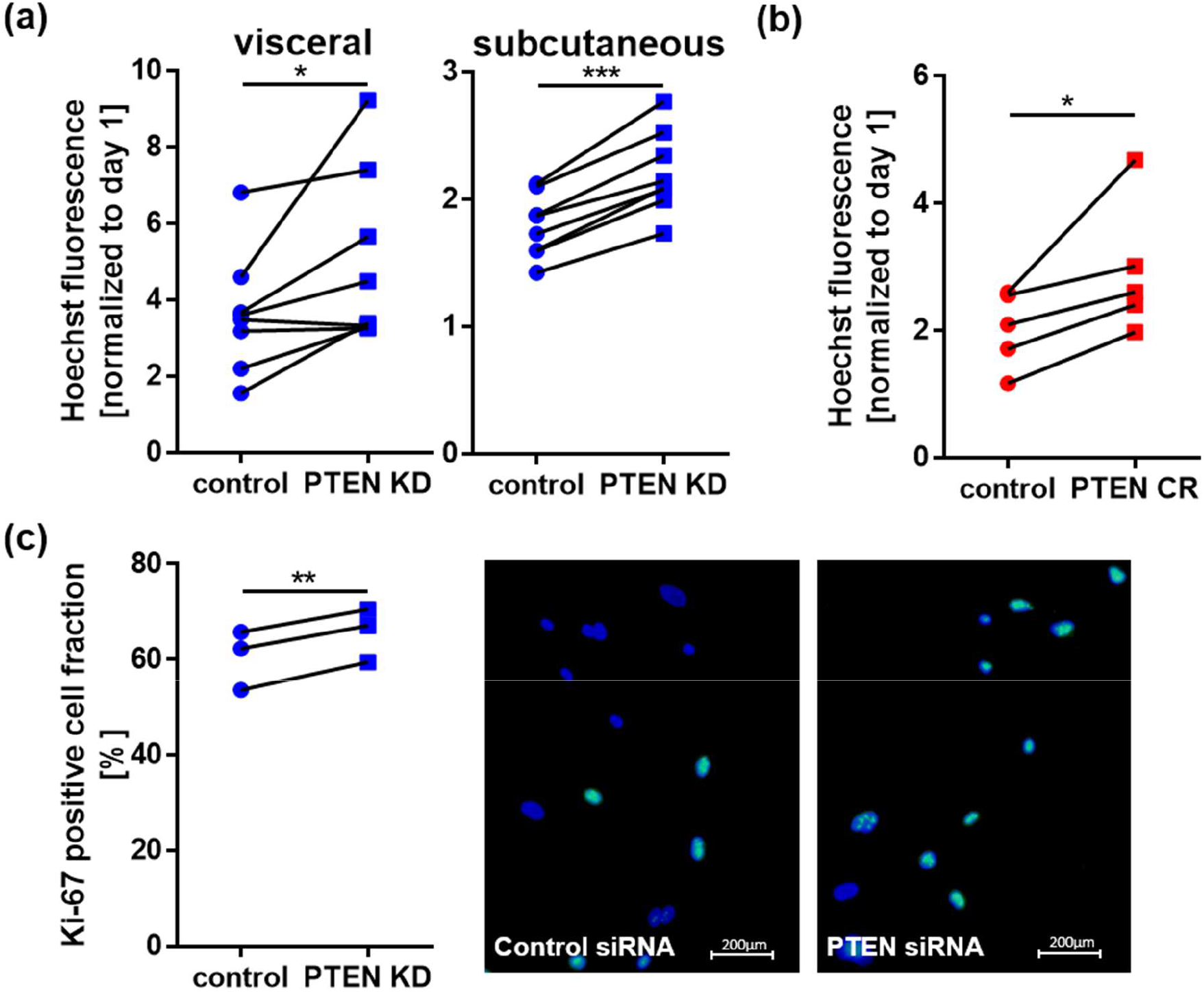
PTEN downregulation enhances proliferation. (a) Hoechst nuclei staining of control and PTEN KD cells 7 days after transfection: Proliferation in PTEN KD cells was increased 1.4 ± 0.2 fold (n=8, p=0.038) in visceral and 1.23 ± 0.02 fold (n=8, p<0.0001) in subcutaneous SVF cells. (b) Hoechst nuclei staining of control and PTEN CR cells 7 days after plating: Proliferation in PTEN CR cells was increased 1.5 ± 0.1 fold (n=5, p=0.039). (c) Hoechst nuclei staining (blue) and Ki-67 immunofluorescence staining (green) in visceral control and PTEN KD cells 1 day after transfection: PTEN KD cells show 1.09 ± 0.01 fold (n=3, p=0.0043) higher fraction of proliferation marker Ki-67 positive cells. Lines between individual data points indicate matched data from single experiments (control vs. respective knockdown). p-values were determined via paired t-test.

### 2.2 PTEN downregulation restored adipogenic potential in high-passage SVF cells

SVF cells lose their capacity for adipocyte differentiation when cultured for several passages (7). In view of the high adipogenic potential observed in PTEN haploinsufficient lipoma cells, we asked whether PTEN downregulation could reverse this process. High-passage SVF cells (> 15 days in culture) which lost their adipocyte differentiation capacity, were transfected with *PTEN* or control siRNA and kept in adipogenic medium for 8 days. Nile Red lipid staining showed a 1.77 ± 0.07 fold (p=0.0026) increase in visceral and 1.44 ± 0.19 fold (p=0.0275) in subcutaneous SVF cells in differentiated cells after PTEN knockdown (Figure 3a). This finding was also supported by an increase in size of 3D spheroids of visceral PTEN KD SVF cells in adipogenic medium (Figure 3b). While the size of control siRNA spheroids remained constant over 12 days, the size of PTEN KD spheroids was increased 1.23 ± 0.03 fold (p<0.001). It was previously shown that the size of the spheroids corresponds to the degree of differentiation (13, 23).

**Figure 3:**
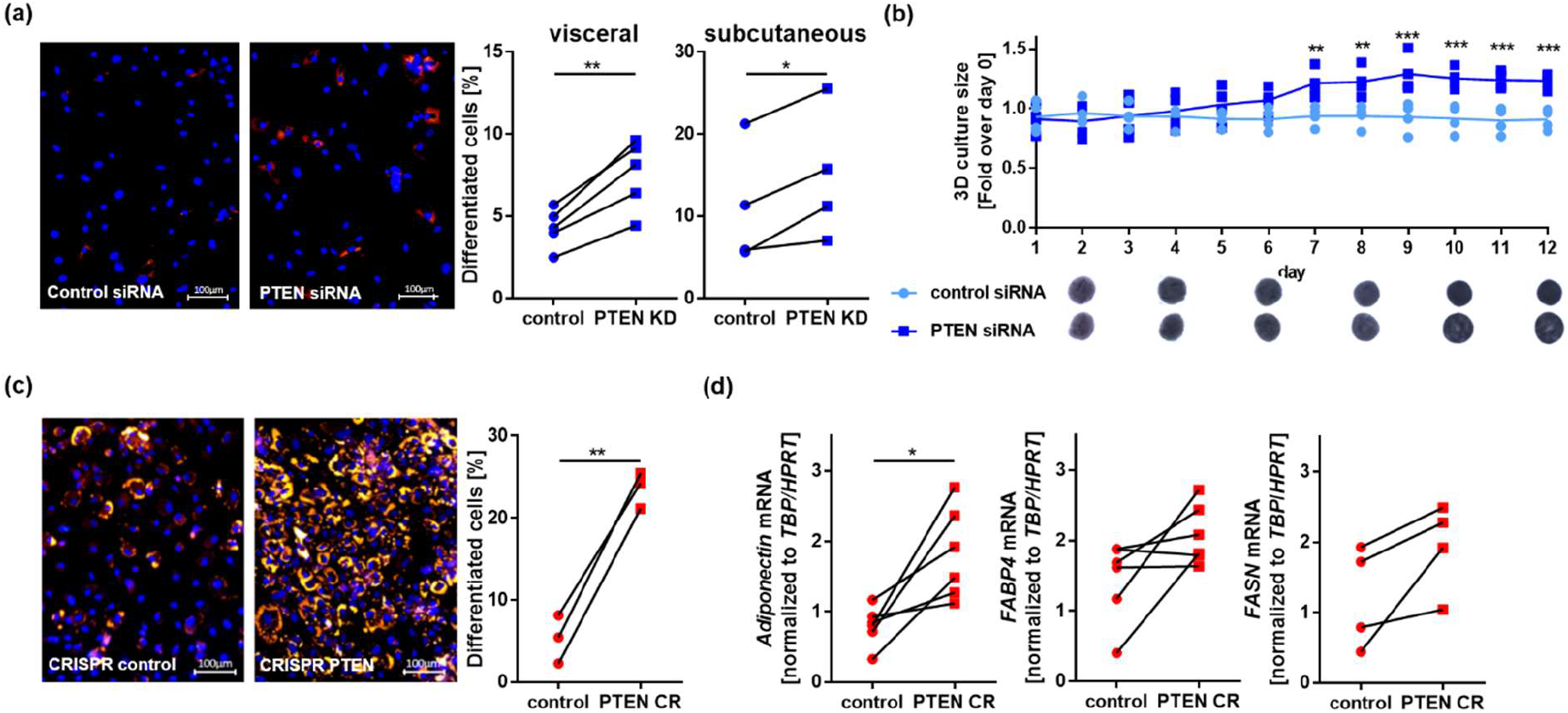
PTEN downregulation enhances adipogenesis. (a) Hoechst nuclei staining (blue) and Nile Red lipid staining (red) in high-passage SVF cells with or without PTEN KD after 8 days in adipogenic medium: The fraction of differentiated cells increased 1.77 ± 0.07 fold (n=5, p=0.0026) in visceral and 1.44 ± 0.19 fold (n=4, p=0.0275) in subcutaneous SVF after PTEN knockdown. (b) 3D culture of visceral control and PTEN KD SVF cells in adipogenic medium for 12 days: The size of control SVF cell spheroids remained constant over 12 days while it was increased 1.23 ± 0.03 fold (n=4, p<0.001) for PTEN KD spheroids (** p<0.01, *** p<0.001). (c) Hoechst nuclei staining (blue) and Nile Red lipid staining (red) in high-passage control or PTEN CR SVF cells after 8 days in adipogenic medium: The fraction of differentiated cells was increased 5.6 ± 1.9 fold (n=3, p=0.004) in CRISPR PTEN knockout cells compared to controls. (d) Expression of the adipocyte markers in control or PTEN CR SVF cells after 8 days in adipogenic medium: *adiponectin* expression was increased in PTEN CR cells 2.6 ± 0.6 fold (normalized to *HPRT* and TBP, n=6, p=0.016), *fatty acid synthase (FASN*) 2.1 ± 0.8 fold (normalized to *HPRT* and TBP, n=4, p=0.075) and *fatty acid binding protein 4 (FABP4*) 1.9 ± 0.6 fold (normalized to *HPRT* and TBP, n=6, p=0.078) compared to controls. Lines between individual data points indicate matched data from single experiments (control vs. respective knockdown). p-values for a), c) and d) were determined via paired t-test, p-values for b) were determined multiple via one-way ANOVA followed by a post hoc Tukey’s multiple comparisons test.

Taking advantage of the stable PTEN downregulation, we observed differentiation capacity of PTEN CR cells 2–6 weeks after transfection. After long term culture only 5 ± 2 % of control cells differentiated into adipocytes during 8 days in adipogenic medium, while 24 ± 1 % of PTEN CR cells differentiated (Figure 3c). This is a 5.6 ± 1.9 fold (p=0.004) increase in differentiation in the PTEN CR cells. Expression of the adipocyte markers *adiponectin* (*ADIPOQ*) (2.6 ± 0.6 fold, p=0.016), *fatty acid synthase* (*FASN*) (2.1 ± 0.8 fold, p=0.075) and *fatty acid binding protein 4* (*FABP4*) (1.9 ± 0.6 fold, p=0.078) was increased in PTEN CR cells compared to controls after 8 days in adipogenic medium, confirming these findings (Figure 3d).

### 2.3 PTEN levels were upregulated during cellular aging

Following the indication of restored differentiation capacity in high passage SVF cells after PTEN downregulation, we investigated whether PTEN levels were regulated during long term culture of SVF cells. We analyzed levels of PTEN protein and AKT phosphorylation (pAKT) in primary visceral SVF cells at different passages. In addition, we detected levels of the nicotinamide adenine dinucleotide (NAD) biosynthesis enzyme nicotinamide phosphoribosyltransferase (NAMPT) which is known to be downregulated during aging (24). PTEN levels rose during long term culture, while the ratio of pAKT (T308) to total AKT levels declined. A decline was also observed in the levels of NAMPT during long term culture (Figure 4a). We observed a reversion in visceral PTEN KD cells, with NAMPT protein 1.6 ± 0.2 fold (p=0.0029) elevated in PTEN KD cells, while levels of the senescence associated cell cycle regulator p21 protein were reduced to 0.33 ± 0.07 fold (p=0.0004) after PTEN KD (Figure 4b). We also found a reduction of *CDKN1A* (*p21*) (to 0.6 ± 0.06 fold, p=0.031) and senescence marker *CDKN2A* (*p16*) (to 0.68 ± 0.05 fold, p=0.014) mRNA in PTEN KD SVF cells (Figure S2A). After long term culture there were 1.58 ± 0.12 fold (p=0.0098) more senescence-associated β-galactosidase (SA-β-gal) positive cells found in the controls compared to PTEN CR cells, supporting a rejuvenating effect of PTEN downregulation (Figure 4c). We detected an upregulation of senescence markers *p21* and *HIPK2* but not *CDKN2B (p15*) in high passage compared to low passage cells from a PHTS patients’ lipoma (LipPD1) (Figure S2B).

**Figure 4:**
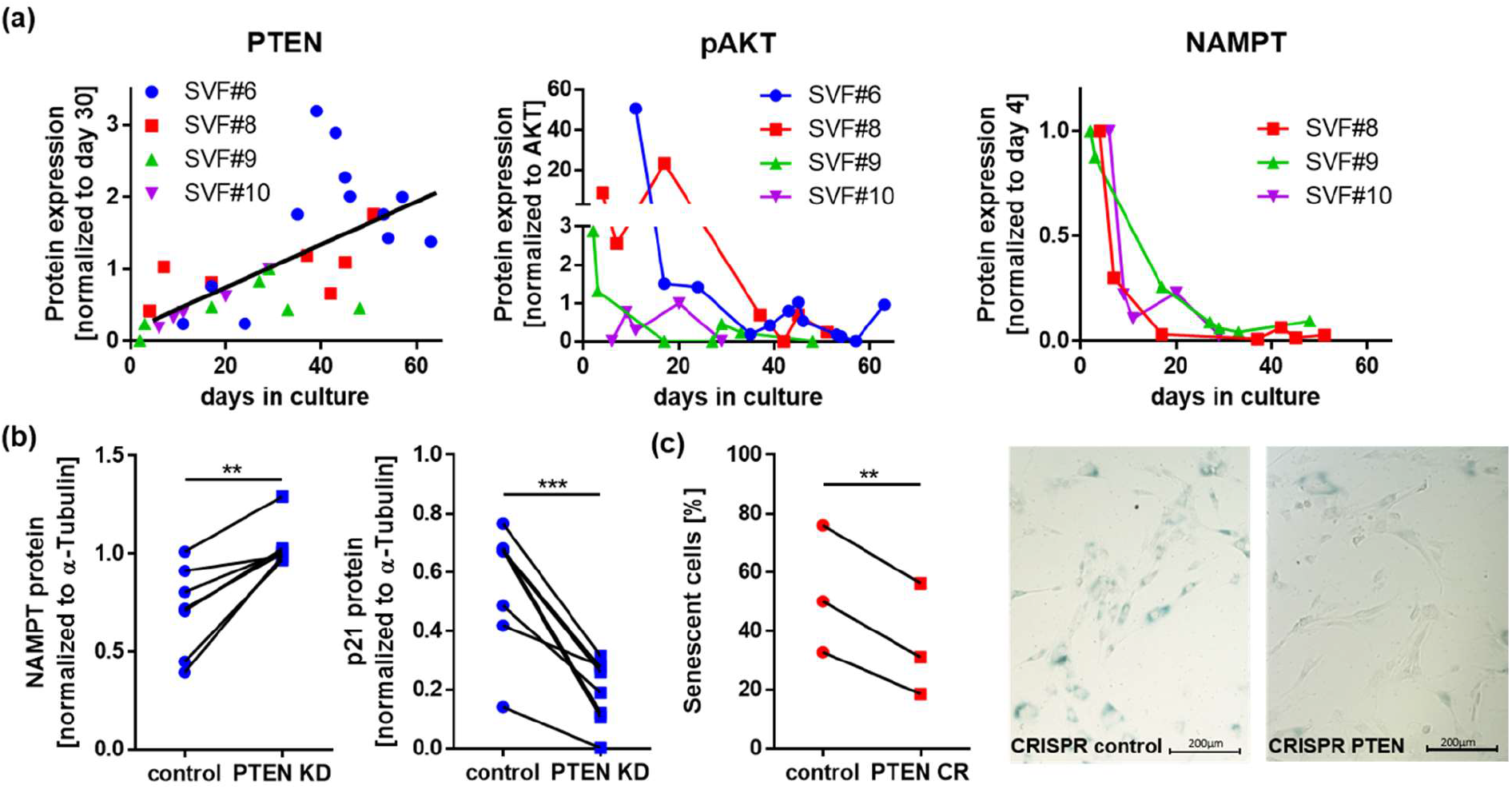
PTEN levels increase during cellular aging. (a) Western blots of primary visceral SVF cells at different days in culture: PTEN levels increased during long term culture (normalized to α-Tubulin, p<0.0001), while the ratio of pAKT (T308) to total AKT (p=0.1) and nicotinamide phosphoribosyltransferase (NAMPT, normalized to α-Tubulin, p=0.0004) decreased during long term culture. (b) Western blots of control and PTEN KD visceral SVF cells: NAMPT protein increased 1.6 ± 0.2 fold (normalized to α-Tubulin, n=7, p=0.0029) while p21 protein decreased to 0.33 ± 0.07 fold (normalized to α-Tubulin, n=8, p=0.0004) in PTEN KD cells. (c) Senescence-associated β-galactosidase (SA-β-gal) assay of control and PTEN CR cells after long term culture: the fraction of SA-β-gal positive cells was 1.58 ± 0.12 fold (n=3, p=0.0098) increased in the controls. Lines between individual data points indicate matched data from single experiments (control vs. respective knockdown). p-values for (a) were determined via linear regression and correlation analysis, p-values for (b) and (c) were determined via paired t-test.

### 2.4 Expression changes after PTEN downregulation

We performed RNA sequencing of PTEN KD and control visceral SVF cells as an untargeted approach to identify genes and pathways regulated in conditions of PTEN downregulation (Tables S1 and S2 available at www.bioinf.uni-leipzig.de/publications/supplements/20-008). We found 1379 differentially expressed genes (adjusted p-value <0.01), of which 170 had a log2(fold change) (log2FC) >1 or <-1 (Figure 5a). More genes were upregulated 60% (829/1379) or 68% (115 of 170) than downregulated, corresponding to the pathway deactivating nature of PTEN (Figure 5b). Among the 50 most significantly differentially expressed genes only two were downregulated in PTEN KD (Figure 5b). Gene set enrichment analysis using StringDB (25) identified 18 significantly enriched Kyoto Encyclopedia of Genes and Genomes (KEGG) pathways (p<0.05) (Figure 5c). The pathway most significantly enriched in PTEN KD compared to control SVF cells was *Cellular Senescence* (hsa04218) with altered gene expression levels for 28 out of 156 genes (p=0.00056). We could confirm the RNA sequencing results for senescence associated genes *p15* and *HIPK2* via qPCR (Figure S2C). Detailed networks of the regulated and other KEGG pathways relevant in the context of metabolism, aging and adipocyte progenitor cells can be found at the supporting information website: www.bioinf.uni-leipzig.de/publications/supplements/20-008.

**Figure 5:**
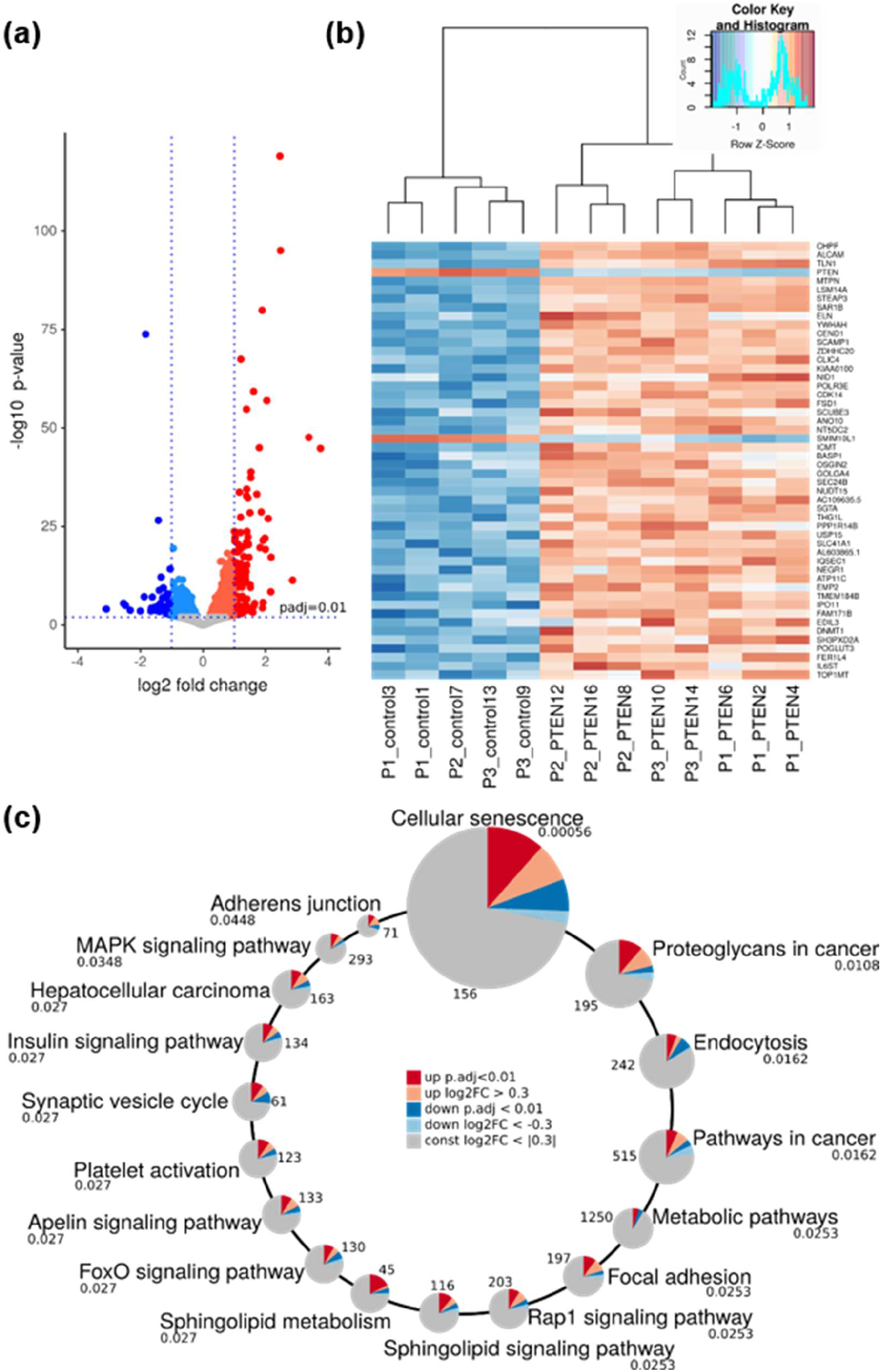
RNA Sequencing of PTEN knockdown and control visceral SVF cells. (a) Volcano plot of differential gene expression in control vs. PTEN KD cells: we found 829 upregulated genes (red) and 550 downregulated genes (blue) (adjusted p<0.01). Of those 115 up- and 55 down-regulated genes had a log2(fold change) (log2FC) >1 or <-1, which means at least duplication or halving of mRNA transcripts. (b) Heatmap of differentially expressed genes in control vs. PTEN KD cells: among the 50 most significantly differentially expressed genes only *PTEN* and *SMIM10L1* were downregulated, while the majority of genes were upregulated. (c) Differentially expressed genes were significantly enriched (adjusted p<0.05) in 18 KEGG-pathways. The circles scale for adjusted p-values, which are given below the pathway name. Numbers next to the circles give the total number of genes assigned to the pathway in KEGG. Dark red and blue slices represent the fractions of significantly up and downregulated genes in the pathways. Light red and blue slices mark additional fractions, where the expression is altered by at least 20% (log2FC >|0.3|), although the expression change was not significant.

### 2.5 Enhanced adipogenesis is mediated through FOXO1 phosphorylation and SREBP1

To identify genes relevant for lipoma development, adipose tissue function and adipogenesis we compared our gene set with two other gene sets; first a study by Le Duc *et al*. (26) comparing lipomas and subcutaneous adipose tissue of the same patient and second a study by Breitfeld *et al*. (27) comparing adipogenesis in murine inguinal and epididymal fat depots. We found an overlap of 36 genes (Table S3), including some known regulators of adipogenesis like *FOXO1* (28) and genes such as *RNF144B* which has no known function in adipose tissue yet. We confirmed the downregulation for these two genes via qPCR (Figure S3A) and analyzed their expression changes during 12 days of adipogenesis in a PHTS patients’ lipoma cells (LipPD1). We found *FOXO1* upregulated and *RNF144B* downregulated during adipocyte development (p<0.001) (Figure 6a). We also investigated the inactivation of FOXO1 through phosphorylation and found that pFOXO1 increased (6.6 ± 3.5 fold, p=0.046) in PTEN KD cells (Figure 6b). The FOXO1 downstream target SREBP1 was upregulated on the protein level (1.5 ± 0.1 fold, p=0.047, Figure 6b) and on the mRNA level (1.6 ± 0.1 fold, p=0.013, Figure S3B) in PTEN KD cells. To investigate whether a reintroduction of the downregulated *FOXO1* and *RNF144B* could attenuate the effects of PTEN downregulation, we overexpressed constitutively active FOXO1 and RNF144B in PTEN CR cells. While both had no consistent effect on proliferation of the PTEN CR cells (Figure S3C), we observed an attenuating effect of constitutively active FOXO1 on adipogenesis (0.73±0.07 fold, p=0,065) while SREBP1 protein levels were reduced (0.49±0.14 fold, p=0.024). RNF144B overexpression had no influence on adipogenesis (Figure S3D).

**Figure 6:**
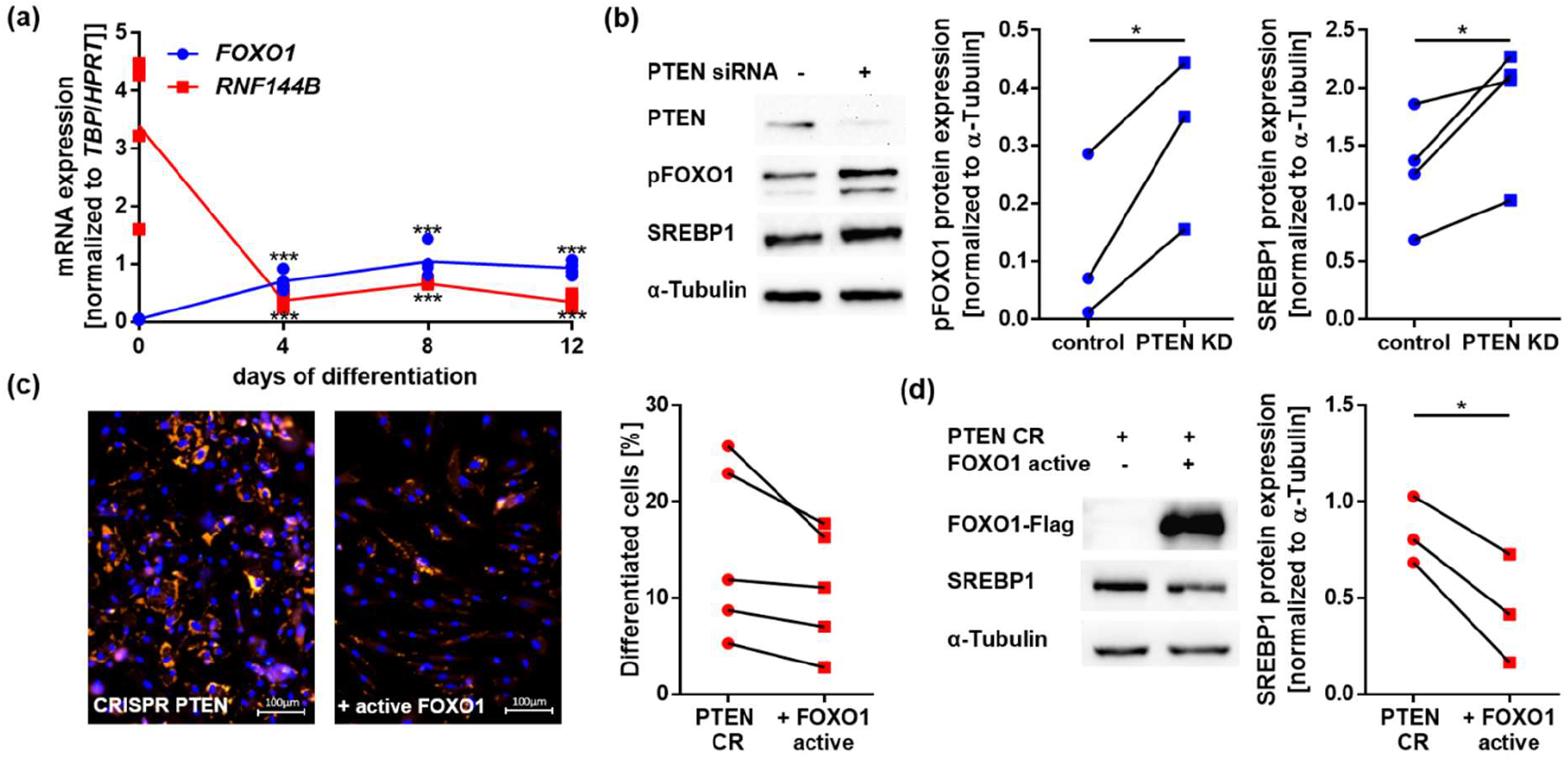
PTEN knockdown induces SREBP1 expression via FOXO1 phosphorylation. (a) Gene expression of FOXO1 and RNF144B during adipogenesis: FOXO1 was upregulated during adipocyte development while RNF144B was downregulated (n=4, *** p<0.001) in lipoma cells from a PHTS patient. (b) Western blots of phosphorylated FOXO1 (pFOXO1) and SREBP1 in control and PTEN KD visceral SVF cells: pFOXO1 was increased 6.6 ± 3.5 fold (n=3, p=0.046) and SREBP1 was increased 1.49 ± 0.13 fold (n=4, p=0.047) after PTEN KD. (c) Hoechst nuclei staining (blue) and Nile Red lipid staining (red) in control and constitutively active FOXO1 overexpressing PTEN CR SVF cells after 8 days in adipogenic medium: The fraction of differentiated cells decreased 0.73±0.07 fold (n=5, p=0.065). (d) Western blots of FOXO1 and SREBP1 in control and constitutively active FOXO1 overexpressing PTEN CR SVF cells: SREBP1 expression decreased 0.49±0.14 fold (n=3, p=0.024) after FOXO1 overexpression. Lines between individual data points indicate matched data from single experiments (control vs. respective knockdown/overexpression). p-values for a) were determined via one-way ANOVA followed by a post hoc Tukey’s multiple comparisons test, p-values for b), c) and d) were determined via paired t-test.

## Discussion

As a well-established tumor suppressor, PTEN is known for its anti-proliferative effects in many cell types. However, not much is known about the role of PTEN in adipocyte progenitors. Both obesity and lipoma formation are characterized by adipose tissue overgrowth. While it is known that in obesity both hypertrophy and hyperplasia contribute to adipose tissue expansion (29), it remains unclear whether these mechanisms also promote lipoma formation. Within this study, we investigated the mechanisms leading to aberrant adipose tissue growth and lipoma formation in patients with heterozygous *PTEN* mutations. A PTEN downregulation in adipocyte progenitor cells of approximately 50 % as observed in lipoma cells of a PHTS patient (13), lead to a several-fold activation of the PI3K downstream targets AKT and ribosomal protein S6. The overactivation of the PI3K pathway enhanced the proliferation of these cells, which is in line with the general cell cycle activating function of the pathway (30). We conclude that PTEN controls adipose tissue growth and inhibits adipocyte progenitor expansion under normal conditions. Conversely, reduction in the phosphatase PTEN may lead to adipose tissue hyperplasia and lipoma formation.

We previously observed that SVF cells obtained from a pediatric PHTS patients’ lipoma retain their ability to differentiate into adipocytes over a prolonged period, while primary SVF cells from healthy donors lose their differentiation capacity after several passages (13). To investigate whether this effect is caused by a reduction in PTEN protein levels, we analyzed differentiation of SVF cells with and without PTEN knockdown. SVF cells that had already lost their capacity to differentiate into adipocytes differentiated again in 2D and 3D culture after siRNA mediated PTEN knockdown. SVF cells with stable CRISPR mediated PTEN downregulation kept their ability to undergo differentiation after long term culture. The SVF cells used within this study were obtained from obese donors and it remains unclear whether this influences the effects seen after PTEN downregulation. We ensured to use SVF cells from non-diabetic donors to avoid altered insulin responses. Enhanced adipogenesis and proliferation was observed both in SVF cells from visceral and subcutaneous adipose tissue depots. In support of the hypothesis that PTEN plays a role in regulating adipogenesis we identified several genes differentially expressed comparing PTEN KD and control cells. We validated two of these, *FOXO1* and *RNF144B*, showing their regulation during adipocyte differentiation. Since adipogenesis can be induced through insulin (5), the stimulating effects of PTEN downregulation on adipogenesis might be mediated through a higher insulin sensitivity. It is known that reduction in PTEN leads to higher insulin sensitivity while body weight increases (31), which could account for these effects. FOXO1 is phosphorylated through insulin signaling via AKT, inhibiting its transcriptional activity (32). We observed increased FOXO1 phosphorylation after PTEN KD and increased expression of the lipogenic transcription factor SREBP1, which is transcriptionally repressed by FOXO1 (33). Overexpression of constitutively active FOXO1 was accompanied by SREBP1 downregulation and led to attenuated adipogenesis in PTEN CRISPR cells. The inhibition of FOXO1 by AKT-mediated phosphorylation may account for the effects on adipogenesis observed after PTEN downregulation. The adipogenic potential of SVF cells from older animals is decreased (4). These age related effects could be coupled to insulin sensitivity, which is also known to decrease with old age (1).

Interestingly, we found that PTEN protein levels are higher in SVF cells after several weeks in culture, connecting replicative age and adipogenic potential, which declines during long term culture. A recent study of tissue specific gene expression in mice during aging by Schaum *et al*. (34) showed a positive correlation of *PTEN* with age in subcutaneous fat depots, while most other tissues (including brown adipose tissue, but not mesenteric adipose tissue) showed a negative correlation. These results indicate a relevance of PTEN in adipose tissue aging, not only *in vitro* but also *in vivo. p16* and *p21* expression as well as SA-β-gal expression were reported to be higher in SVF cells from older donors (35). To test whether there is a direct effect of PTEN on cellular senescence, we compared *p16* and *p21* expression with and without PTEN knockdown and found a reduction in knockdown cells. We observed less SA-β-gal positive cells in SVF cells with CRISPR mediated PTEN downregulation. These results provide evidence for a connection between PTEN expression and cellular senescence, though replicative aging processes are not completely blocked in PTEN haploinsufficient cells, as seen in comparison of senescence markers in low and high passage LipPD1 cells. In contrast to pathway activation through PTEN downregulation, PI3K inhibition with alpelisib induced senescence in PTEN haploinsufficient lipoma cells (9) and PTEN overexpression was shown to induce G1 arrest in cancer cells mediated through AKT inhibition (36). We found *Cellular Senescence* to be the most significantly enriched KEGG-pathway in RNA sequencing of PTEN knockdown vs. control SVF cells, with senescence related genes like *CDKN2B* (37) and *HIPK2* (38) downregulated. A relation between PI3K/AKT pathway and aging was also reported in other stem cell types: mTOR complex 1 (mTORC1) activity in intestinal stem cells declines with aging (39). Age related upregulation of the upstream antagonist PTEN might be responsible for this decline in downstream PI3K signaling pathway activity and could also account for the age related decline in MSCs proliferation rate (35). On the other hand, there is strong evidence that inhibition of the insulin signaling pathway with a special focus on mTORC1 increases longevity. This makes sense in the context of insulin induced metabolic activity and increased reactive oxygen species (ROS) production (40). Some authors suggest an over proliferation and exhaustion of adipose progenitors in conditions of nutrient abundance associated with telomere shortening (41). Interestingly, Schumacher *et al*. found similarities in differential gene expression between mouse models of delayed and premature aging compared to wild-type mice, with downregulation of IGF-1 related pathways as a common feature (42). Varying PTEN expression might be a cellular mechanism to maintain a balance between senescence and over proliferation, ensuring normal adipose tissue homeostasis. In disease states PTEN downregulation leads to excessive adipose tissue growth and distribution abnormalities, while PTEN upregulation might contribute to senescence and dysfunctional adipose tissue in older individuals.

## Experimental procedures

### 4.1 Cell culture and adipocyte differentiation

We used cells of the stromal vascular fraction isolated from visceral or subcutaneous adipose tissue of healthy donors, resected during bariatric surgery as well as PTEN-haploinsufficient lipoma cells (LipPD1) from a pediatric PHTS patient (13). The study was approved by the Leipzig University ethics committee (ethical approval: no. 425-12-171220) and has been performed in accordance with the principles laid down in the 1964 Declaration of Helsinki and its later amendments. All adipose tissue and lipoma tissue donors gave written informed consent to participate in the study. Isolation and culture methods were described previously (16, 43). Supplementary Table S4 contains a list of the primary cell cultures used. To characterize the cell populations found in these cultures, cell surface markers from SVF and LipPD1 cells cryopreserved at different passages were determined via flow cytometry analysis. Cells were thawed and resuspended in PBS supplemented with 0.5% BSA and 2mM EDTA (pH 7.4). The cells were incubated 30 minutes at 4°C, with fluorescently labeled monoclonal anti-human antibodies specific for CD8-PerCP, CD31-V450 (BD Biosciences), MSCA1-PE, CD271-APC, CD14-PeVio770, CD34-FITC (Miltenyi Biotec), and CD45-BV510 (BioLegend) (Table S5) (44). After washing steps, the cells were analyzed by flow cytometry using a FACS Aria II SORP flow cytometer and Diva Pro software (BD Biosciences) at the Core Unit Fluorescence Technologies of the University of Leipzig. Results from the analysis are presented in Table S6.

For differentiation 15,000 cells/96-well or 120,000/12-well were plated in culture medium. Medium was changed to differentiation medium 24 h later (day 0) (DMEM/F12 containing 8 mg/ml D-biotin, 10 mg/ml D-pantothenic acid, 2 μM rosiglitazone, 25 nM dexamethasone, 0.5 mM methylisobuthylxantine, 0.1 μM cortisol, 0.01 mg/ml apotransferrin, 0.2 nM triiodotyronin, and 20 nM human insulin (45) and cells were kept in differentiation medium for 4, 8 or 12 days.

A modified method according to Klingelhutz *et al*. (23) was used for scaffold-free 3D cultures of SVF cells. After transfection 10,000 cells per well were seeded into low attachment 96-well microplates (PS, U-bottom, clear, cellstar^®^, cell-repellent surface, Greiner Bio-One) in 100 μl culture medium. After one day medium was changed to differentiation medium and spheroids were kept in differentiation medium for 12 days. Half of the medium was replaced every second day. Microscope images were taken daily using the EVOS FL Auto 2 Cell Imaging System (Invitrogen; Thermo Fisher Scientifc). Image analysis to determine the spheroid size was performed using ImageJ (46).

### 4.2 PTEN siRNA transfection

One day prior to transfection, cells were plated at a density of about 1,300 cells/cm^2^ to ensure optimal growth. For transfection we used the Neon Transfection System 100 μl Kit (Invitrogen; Thermo Fisher Scientific, Inc.). Cells were harvested via trypsination and washed with DPBS. Either control siRNA (Silencer^™^ Negative Control No. 1 siRNA, Ambion, Thermo Fisher Scientific, Inc.) or a combination of *PTEN* siRNA (s325 and s326, both Ambion, Thermo Fisher Scientific, Inc.) was added to the cell pellets (final concentration of 10 nM in culture medium after transfection). Pellets were resuspended in 100 μl R-buffer for transfection and electroporated in Neon 100 μl tips at 1,300 V, 2 pulses, 20 ms. After electroporation cells were transferred to prewarmed culture medium and distributed to multiwell tissue culture plates for functional assays.

### 4.3 CRISPR/Cas *PTEN* knockout

For stable knockout of *PTEN* in SVF cells, we used the CRISPR/Cas9 genome editing technique in a reverse transfection of guideRNA/Cas9 nickase ribonucleoproteins (RNPs). Cas9 nickase was chosen to avoid off-target effects. If not otherwise stated, reagents were purchased from Integrated DNA Technologies (Alt-R CRISPR-Cas9 system). Transfections were performed according to the manufacturers’ protocol with a combination of two different crRNA:tracrRNA guides and Alt-R S.p. Cas9 D10A Nickase 3 NLS (# 1078729). We tested combinations of four different crRNAs (Table S7) for their editing efficiency via T7EI assay (Alt-R Genome Editing Detection Kit, # 1075931). For further assays we used the most efficient guide combination (#3 and #4). RNPs of Cas9 nickase and Alt-R CRISPR-Cas9 Negative Control crRNA #1 (#1072544) were used for controls.10,000 SVF cells per well were transfected in a 48-well format with a final concentration of 10 nM crRNA:tracrRNA:Cas9 nickase RNPs. We used a total volume of 300 μl per well with 2.4 μl Lipofectamin RNAiMAX (Thermo Fisher Scientific, #13778075). Efficiency was determined 2 days post transfection via T7EI assay.

### 4.4 FOXO1 and RNF144B overexpression

Constitutively active FOXO1 and RNF144B were overexpressed in PTEN CR and LipPD1 cells. One day prior to transfection, cells were plated at a density of about 1,300 cells/cm^2^ to ensure optimal growth. For transfection we used the Neon Transfection System 100 μl Kit (Invitrogen; Thermo Fisher Scientific, Inc.). Cells were harvested via trypsination and washed with DPBS. Either a control vector (c-Flag pcDNA3 was a gift from Stephen Smale (Addgene plasmid #20011; http://n2t.net/addgene:20011; RRID:Addgene_20011(47)), a vector for overexpression of constitutively active FOXO1 with mutated phosphorylation site (pcDNA3 Flag FKHR AAA mutant was a gift from Kunliang Guan (Addgene plasmid #13508; http://n2t.net/addgene:13508; RRID:Addgene_13508) (32)) or a vector for RNF144B overexpression (RNF144B pcDNA3.1+ C-(K)-DYK #OHu07981D, GenScript) was added to the cell pellets (final concentration of 1 μg/ml for FOXO1 and 2 μg/ml for RNF144B in culture medium after transfection). Pellets were resuspended in 100 μl R-buffer for transfection and electroporated in Neon 100 μl tips at 1,300 V, 2 pulses, 20 ms. After electroporation cells were transferred to prewarmed culture medium and distributed to multiwell tissue culture plates for functional assays.

### 4.5 Lipid staining

For lipid staining cells were fixed in 4% PFA and washed with DPBS. Afterwards cells were co-stained with the fluorescent dyes Nile Red (0.5 μg/ml, Sigma) and Hoechst 33342 (1 μg/ml, Sigma) for 5 min in DPBS. Microscope images were taken with the EVOS FL Auto 2 Cell Imaging System (Invitrogen; Thermo Fisher Scientific) and cell counting was performed with the Celleste Image Analysis Software (Thermo Fisher Scientific).

### 4.6 Proliferation assay

For proliferation assays cells were seeded at a density of 2,000 cells/well on 96-well plates. Growth medium was replaced every 72 h. Cells were fixed at day 1 and day 7 after transfection and kept in DPBS at 4 °C after fixation. Hoechst 33342 (Sigma) was used to stain nuclei for 5 min at a concentration of 1 μg/ml in DPBS. Hoechst fluorescence was detected at 455 nm and compared for day 1 and day 7.

### 4.7 Western blot analysis

For Western blot analysis transfected cells were seeded at a density of 10,000 cells/cm2 in culture medium. Cell culture medium was replaced by serum free medium one day after transfection and cells were harvested on the next day. For Western blot analysis of long term cultures, cells were trypsinized during normal culture and 100,000 cells were frozen as a pellet. Proteins were extracted and immunoblotting was performed as described elsewhere (16). We used 10 μg protein per lane and incubated with primary antibodies (Cell Signaling Technology, CST) and secondary antibodies (Dako; Agilent Technologies) according to Table S8. α-Tubulin was used as loading control. Densitometric analysis was performed using ImageJ (46) and images of exposed films and their analysis are provided in Figure S4 (PTEN KD), Figure S5 (PTEN CR) and Figure S6 (long term culture).

### 4.8 Immunofluorescence staining

For Ki-67 immunofluorescence staining transfected cells were seeded at a density of 2,000 cells/well on 96-well plates and fixed with 4 % PFA after 24 h. Cells were permeabilized and blocked in IF-buffer (DPBS + 5 % BSA + 0.3 % Tween20) for 1 h at room temperature (RT) and stained with Ki-67 primary antibody over night at 4 °C (Supplementary Table S8). Cells were washed three times 5 min with IF-buffer and incubated with secondary Alexa Fluor 488 antibody (Table S8) 2 h at RT in the dark. Afterwards cells were washed one time with IF-buffer, then incubated 5 min with 1 μg/ml Hoechst 33342 in DPBS and washed one time with DPBS. Microscope images were taken in DPBS using the EVOS FL Auto 2 Cell Imaging System (Invitrogen; Thermo Fisher Scientific) and cell counting was performed with the Celleste Image Analysis Software (Thermo Fisher Scientific).

### 4.9 Reverse transcription quantitative PCR (RT-qPCR)

We plated 6,000 cells/cm^2^ for RT-qPCR of undifferentiated cells or 45,000 cells/cm2 for differentiated cells. RNA extraction, reverse transcription and qPCR were performed as previously described (43). Supplementary Table S9 contains a list of primers used for qPCR assays. Results were normalized to the housekeepers *hypoxanthine phosphoribosyltransferase* (*HPRT*) and *TATA box binding protein (TBP*). We performed probe based assays using the Takyon^™^ Low Rox Probe MasterMix dTTP Blue (Eurogentec) or SYBR green assays using the Takyon^™^ Low Rox SYBR^®^ MasterMix dTTP Blue (Eurogentec).

### 4.10 RNA sequencing

After PTEN knockdown we plated 6,000 cells/cm2 in culture medium. Medium was replaced by serum free medium 1 day after transfection and cells were harvested 24 h afterwards. RNA was extracted as for RT-qPCRs. 100 ng of total RNA was used for library synthesis with the NEBNext Ultra II Directional RNA Library Preparation kit (New England Biolabs) according to the protocol of the manufacturer. The barcoded libraries were purified and quantified using the Library Quantification Kit - Illumina/Universal (KAPA Biosystems) on a TaqMan 7500 RealTime PCR System. A pool of up to 10 libraries was used for cluster generation at a concentration of 10 nM. Sequencing of 2×150 bp was performed with a NextSeq 550 sequencer (Illumina) at the sequencing core facility of the Faculty of Medicine (Leipzig University) according to the instructions of the manufacturer.

#### 4.10.1 Read preprocessing and mapping

Sequencing reads from 18 fastq files (triplicates of PTEN KD and controls from 3 donors) were mapped to the human reference genome (hg38 downloaded from UCSC/Ensembl (48)). Paired-end RNA sequencing data were processed within the Galaxy platform (49). The raw sequencing reads (in fastq-format) were loaded to Galaxy instance and quality was inspected with fastqc (50). One sample (PTEN KD and the respective control) had to be removed from further analysis due to low quality. Reads were trimmed with TrimGalore! (51) and sufficient quality was ensured by a second quality check with fastqc. Subsequently, the reads were mapped to the human genome (hg38) with segemehl (52) and annotated with gencode.v29.

#### 4.10.2 Differential gene expression and gene set enrichment

For differential gene expression analysis with DESeq2 (53) only genes with at least ten mapped reads in at least half of the samples were kept, reducing the analysis from 58,721 to 32,660 genes. After inspection of *PTEN* expression levels in all samples, we decided to exclude controls with only moderate *PTEN* expression levels to ensure clear distinguishability between high *PTEN* expression levels and *PTEN* knockdown. Thus controls 5, 11, and 15 were excluded from further analyses (Figure S7A). Afterwards, we ran DESeq2 to identify differentially expressed genes. A principle component analysis identified the different donors are the main source of variance, thus we corrected for the donor (batch) effect (Figure S5B), leading to better clustering of PTEN KD vs. control. We applied LFC shrinkage to reduce effect sizes of overall low expressed genes (shrinkage estimator ashr) (54).

Gene set enrichment analysis was performed in STRING (string-db.org) (25). As input the 1379 significantly differentially expressed genes were used. The results were visualized using R custom scripts and the KEGG pathway visualization package pathview (55).

### 4.11 Statistical analysis

Means of at least three independent experiments were statistically analyzed using GraphPad Prism 6 software (GraphPad Software Inc.). For comparison of control and conditions of PTEN downregulation (PTEN KD or PTEN CR), means of independent experiments were compared via paired Student’s t-test (comparison of each control transfection with the respective PTEN siRNA/PTEN CRISPR transfection). For multiple comparisons we used one- or two-way analysis of variance (ANOVA) followed by a post hoc Tukey’s multiple comparisons test (one-way ANOVA) or Dunnett’s multiple comparisons test (two-way ANOVA). To determine the significance of fold changes, we used one-sample t-tests of the log(fold change) and compared to the hypothetical value 0 (56). Results were indicated as mean ± standard error of means (SEM).

## Data availability statement

All data are contained within the article and in the supporting information. Gene lists and other supporting information are also available at http://www.bioinf.uni-leipzig.de/publications/supplements/20-008.

## Supporting information

This article contains supporting information.

## Acknowledgments

We thank Prof. Dr. Arne Dietrich for giving us access to and all donors for providing adipose tissue samples. We thank Sandy Richter and Anja Barnikol-Oettler for the technical support in the laboratory. This work was supported by the Core Unit Fluorescence Technologies and sequencing core facility of the Faculty of Medicine of the Leipzig University. We thank Janine Obendorf and Sören Pietsch for their help with designing ImageJ plugins for 3D culture image analysis.

## Authors’ contributions

AG, WK and ASK designed the study. ASK, AG, MN, MH and JL performed the experimental work. SK performed the analysis and presentation of RNA sequencing results. DLD and JB provided their RNA sequencing results. ASK prepared the manuscript draft. All authors reviewed and edited the manuscript. Project administration, supervision and funding acquisition: AG, WK, MP, PFS, KL, AK and PK.

## Funding and additional information

The project was supported by the German Research Foundation (209933838 - SFB 1052/Deutsche Forschungsgemeinschaft) and Mitteldeutsche Kinderkrebsforschung Stiftung für Forschung und Heilung. DLD is funded through “Clinician Scientist Program, Medizinische Fakultät der Universität Leipzig.” MH was supported through the ESPE Early Career Scientific Development Grant. KL and AK were supported by the Federal Ministry of Education and Research (BMBF), Germany, (FKZ: 01EO1501, IFB AdiposityDiseases). PFS is also affiliated with the Institute of Theoretical Chemistry of the University of Vienna, Austria, the National University of Colombia, and the Santa Fe Institute. We acknowledge support from Leipzig University for Open Access Publishing.

## Supporting information figure legends

Complete supporting information is available at http://www.bioinf.uni-leipzig.de/publications/supplements/20-008.

**Figure S1:**
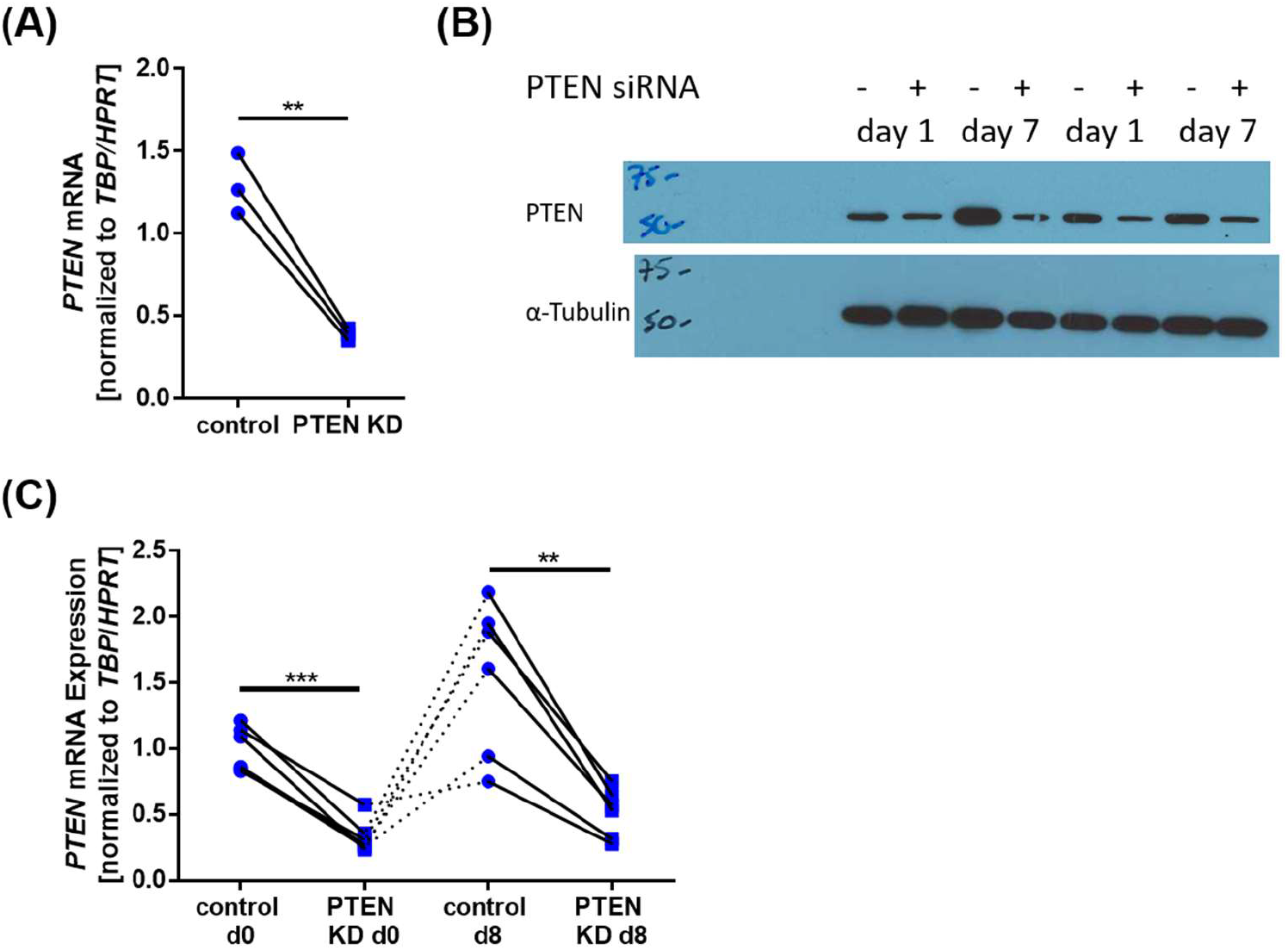
PTEN expression was reduced in control and PTEN KD SVF cells. (A) PTEN mRNA expression in subcutaneous PTEN KD SVF cells was downregulated 0.3 ± 0.01 fold (n=3, p=0.009) compared to controls. (B) Western blots of visceral PTEN KD/control SVF cells: PTEN knockdown was stable for 7 days after siRNA transfection. (C) *PTEN* mRNA expression in visceral PTEN KD/ SVF cells was downregulated upon induction of differentiation (d0, one day after transfection) and after eight days of adipose differentiation (d8) compared to controls. Lines between individual data points indicate matched data from single experiments (control vs. respective knockdown). p-values were determined via paired t-test (** p<0.001 *** p<0.001).

**Figure S2:**
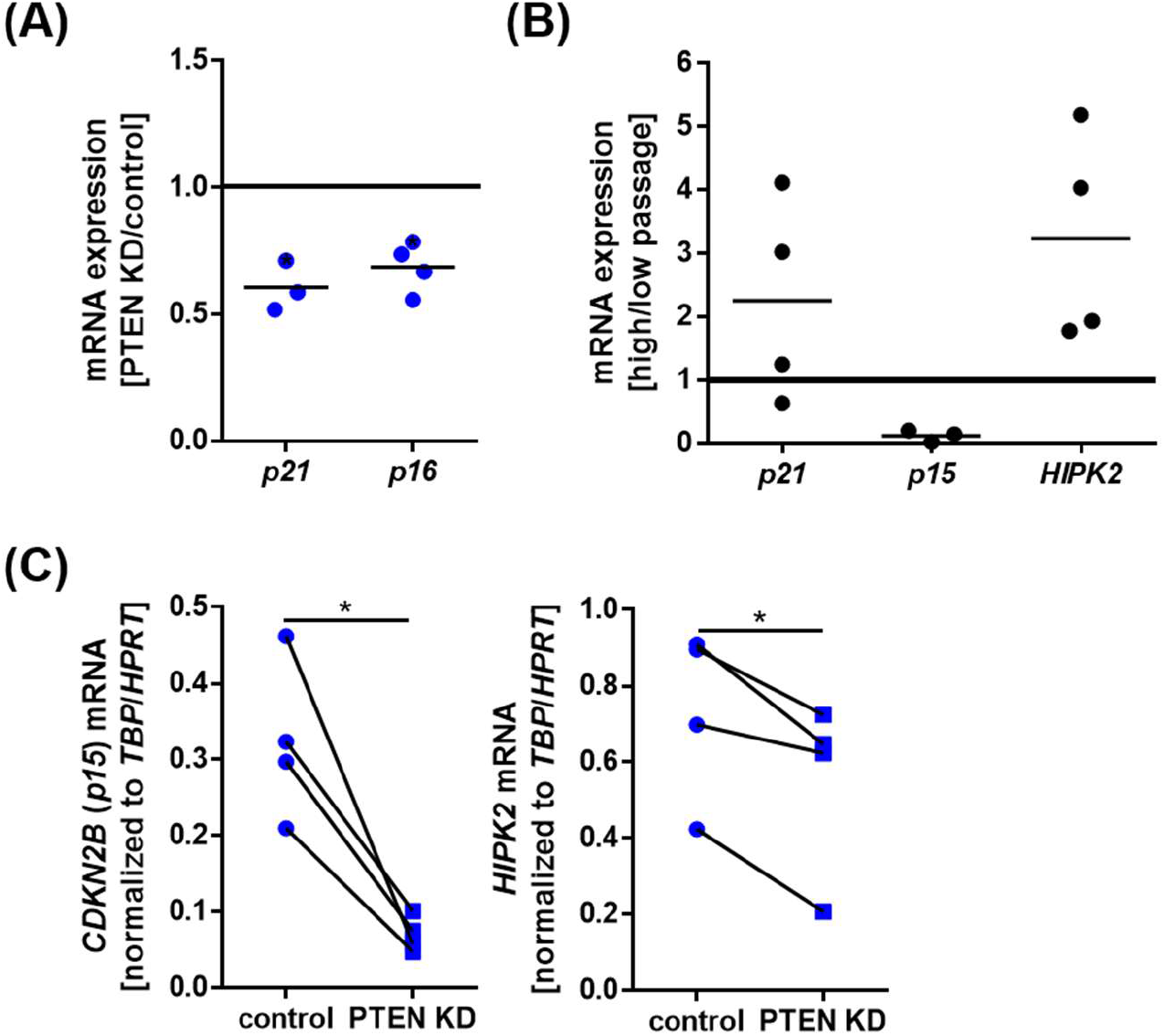
mRNA expression of senescence marker was reduced after PTEN KD. (A) Senescence marker qPCRs of PTEN KD/control visceral SVF cells: On the mRNA level we found a reduction of *CDKN1A* (*p21*) (to 0.6 ± 0.06 fold, n=3, p=0.031) and senescence marker *CDKN2A* (*p16*) (to 0.68 ± 0.05 fold, n=4, p=0.014). (B) Senescence marker qPCRs comparing low passage vs. high passage LipPD1 cells: On the mRNA level we found *p21* elevated 2.3 ± 0.8 fold, (n=4, p=0.27), *CDKN2B* (*p15*) reduced to 0.02 ± 0.05 fold (n=3, p=0.07) and HIPK2 elevated 3.2 ± 0.8 fold (n=4, p=0,028). (C) qPCRs to confirm results from RNA-sequencing: *p15* (to 0.23 ± 0.04 fold, n=4, p=0.017) and *HIPK2* (to 0.73 ± 0.09 fold, n=4, p=0.02) were downregulated after PTEN KD in visceral SVF cells. Lines between individual data points indicate matched data from single experiments (control vs. respective knockdown). p-values for (A) and (B) were determined via one-sample t-test of the log(fold change), p-values for (C) were determined via paired t-test.

**Figure S3:**
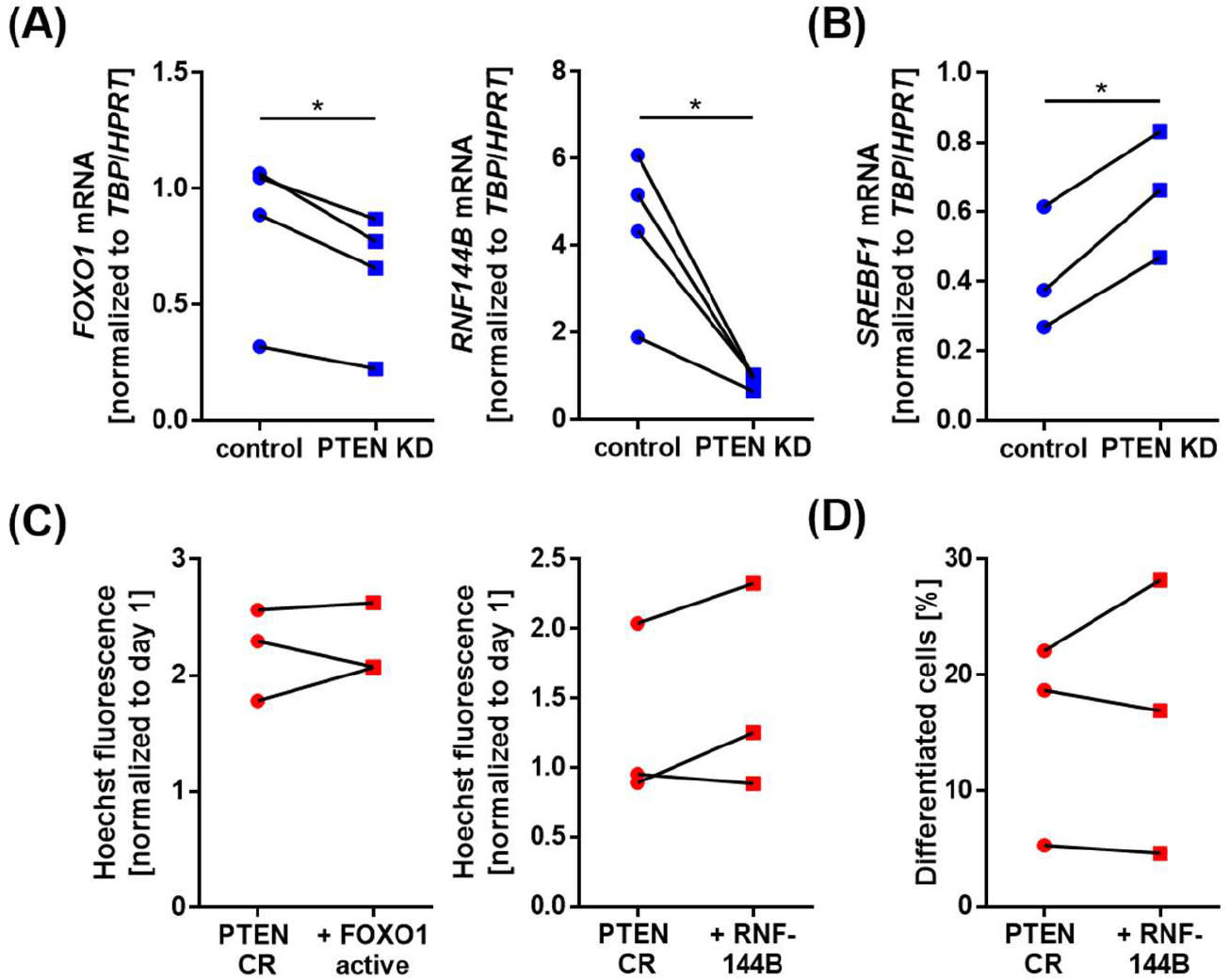
FOXO1 and RNF144B are downregulated in PTEN KD cells. (A) qPCRs to confirm results from RNA-sequencing: *FOXO1* (to 0.75 ± 0.03 fold, n=4, p=0.018) and *RNF144B* (to 0.23 ± 0.04 fold, n=4, p=0.025) were downregulated after PTEN KD in visceral SVF cells. (B) *SREBF1* was upregulated on the mRNA level (1.6 ± 0.1 fold, n=3, p=0.013) in visceral PTEN KD cells. (C) Neither overexpression of constitutively active FOXO1 nor RNF144B in PTEN CR cells had effects on proliferation (as determined via Hoechst assay 7 days post transfection normalized to day 1 after transfection). (D) RNF144B overexpression had no influence on adipogenesis of PTEN CR cells. Lines between individual data points indicate matched data from single experiments (control vs. respective knockdown or overexpression). p-values were determined via paired t-test.

**Figure S4**: Western blots of PTEN KD/control SVF cells: exposed films and densitometric analyses of PTEN (normalized to α-Tubulin), pAKT (T308) (normalized to total AKT), pS6 (Ser235/236) (normalized to α-Tubulin), NAMPT(normalized to α-Tubulin), p21 (normalized to α-Tubulin), pFOXO1 (normalized to α-Tubulin) and SREBP1 (normalized to α-Tubulin).

**Figure S5**: Western blots of PTEN CR/control SVF cells: exposed films and densitometric analyses of PTEN (normalized to α-Tubulin), pAKT (T308) (normalized to total AKT), pS6 (Ser235/236) (normalized to α-Tubulin), FOXO1 (FLAG antibody, normalized to α-Tubulin) and SREBP1 (normalized to α-Tubulin).

**Figure S6**: Western blots of four SVF cell long term cultures: exposed films and densitometric analyses of PTEN (normalized to α-Tubulin), pAKT (T308) (normalized to total AKT) and pS6 (Ser235/236) (normalized to α-Tubulin).

**Figure S7:**
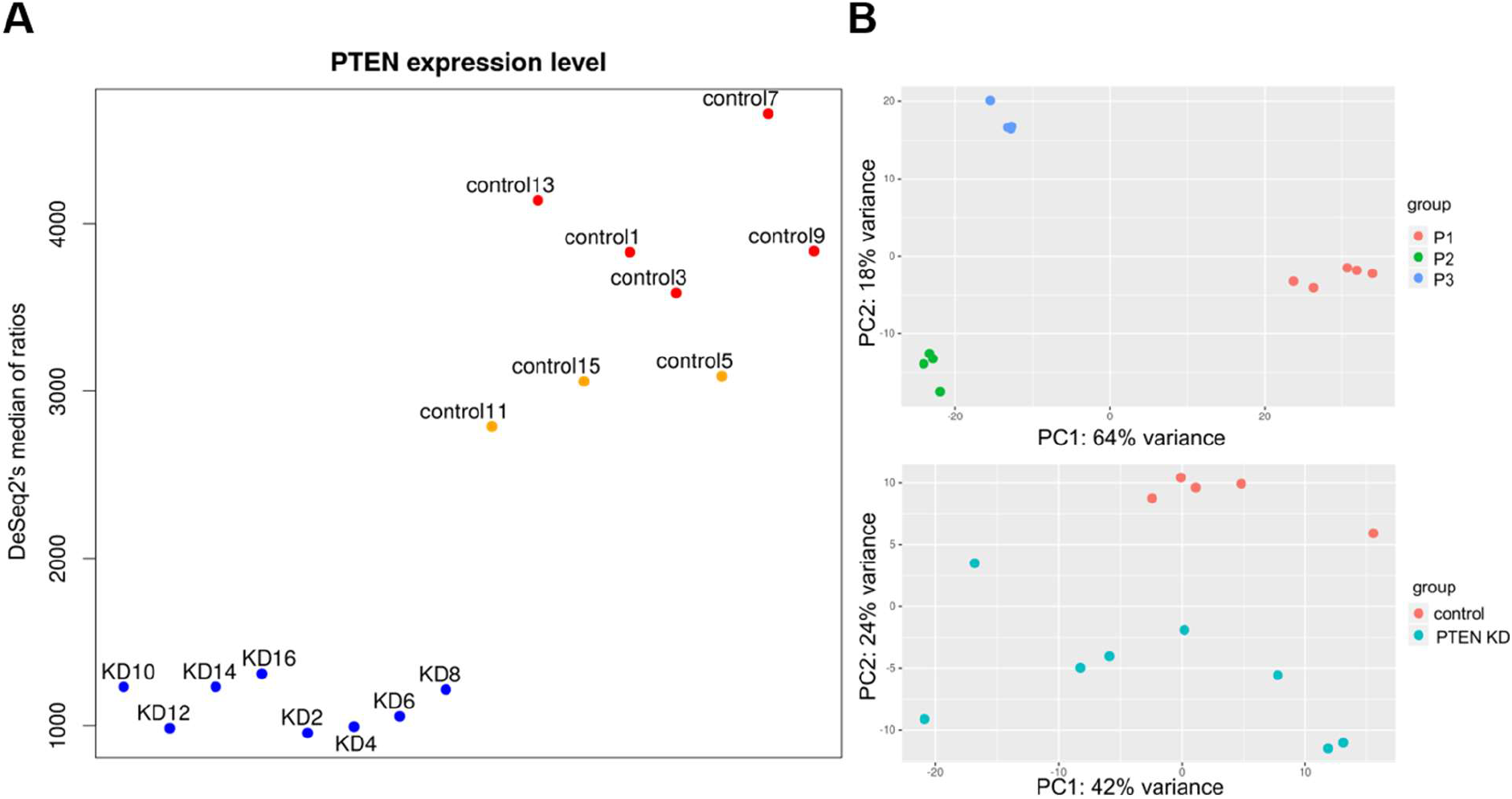
(A) Normalized read counts mapping to *PTEN* mRNA in the investigated samples. (B) PCA of samples clearly cluster according patients (upper panel), after removing the variance caused by the individual cell donors the second principal component (PC2) segregates the samples into control vs. PTEN (lower panel).

## References

1. Kirkland, J. L., Tchkonia, T., Pirtskhalava, T., Han, J., and Karagiannides, I. (2002) Adipogenesis and aging: Does aging make fat go MAD? Experimental Gerontology 37, 757–767 10.1016/S0531-5565(02)00014-1

2. Tchkonia, T., Morbeck, D. E., Zglinicki, T. von, van Deursen, J., Lustgarten, J., Scrable, H., Khosla, S., Jensen, M. D., and Kirkland, J. L. (2010) Fat tissue, aging, and cellular senescence. Aging cell 9, 667–684 10.1111/j.1474-9726.2010.00608.x PMID 20701600

3. Yki-Jarvinen, H., Kiviluoto, T., and Nikkila, E. A. (1986) Insulin binding and action in adipocytes in vitro in relation to insulin action in vivo in young and middle-aged subjects. Acta endocrinologica 113, 88–92 10.1530/acta.0.1130088 PMID 3532658

4. Cartwright, M. J., Tchkonia, T., and Kirkland, J. L. (2007) Aging in adipocytes: Potential impact of inherent, depot-specific mechanisms. Experimental Gerontology 42, 463–471 10.1016/j.exger.2007.03.003 PMID 17507194

5. Cignarelli, A., Genchi, V. A., Perrini, S., Natalicchio, A., Laviola, L., and Giorgino, F. (2019) Insulin and Insulin Receptors in Adipose Tissue Development. International journal of molecular sciences 20 10.3390/ijms20030759 PMID 30754657

6. Hauner, H., Entenmann, G., Wabitsch, M., Gaillard, D., Ailhaud, G., Negrel, R., and Pfeiffer, E. F. (1989) Promoting effect of glucocorticoids on the differentiation of human adipocyte precursor cells cultured in a chemically defined medium. The Journal of clinical investigation 84, 1663–1670 10.1172/JCI114345 PMID 2681273

7. Wabitsch, M., Brenner, R. E., Melzner, I., Braun, M., Möller, P., Heinze, E., Debatin, K. M., and Hauner, H. (2001) Characterization of a human preadipocyte cell strain with high capacity for adipose differentiation. International journal of obesity and related metabolic disorders: journal of the International Association for the Study of Obesity 25, 8–15 10.1038/sj.ijo.0801520 PMID 11244452

8. Bell, A., Grunder, L., and Sorisky, A. (2000) Rapamycin inhibits human adipocyte differentiation in primary culture. Obesity research 8, 249–254 10.1038/oby.2000.29 PMID 10832768

9. Kirstein, A. S., Augustin, A., Penke, M., Cea, M., Körner, A., Kiess, W., and Garten, A. (2019) The Novel Phosphatidylinositol-3-Kinase (PI3K) Inhibitor Alpelisib Effectively Inhibits Growth of PTEN-Haploinsufficient Lipoma Cells. Cancers 11 10.3390/cancers11101586 PMID 31627436

10. Blüher, M., Kahn, B. B., and Kahn, C. R. (2003) Extended longevity in mice lacking the insulin receptor in adipose tissue. Science (New York, N.Y.) 299, 572–574 10.1126/science.1078223 PMID 12543978

11. Bartke, A. (2008) Impact of reduced insulin-like growth factor-1/insulin signaling on aging in mammals: Novel findings. Aging cell 7, 285–290 10.1111/j.1474-9726.2008.00387.x PMID 18346217

12. Katic, M., Kennedy, A. R., Leykin, I., Norris, A., McGettrick, A., Gesta, S., Russell, S. J., Bluher, M., Maratos-Flier, E., and Kahn, C. R. (2007) Mitochondrial gene expression and increased oxidative metabolism: Role in increased lifespan of fat-specific insulin receptor knock-out mice. Aging cell 6, 827–839 10.1111/j.1474-9726.2007.00346.x PMID 18001293

13. Kässner, F., Kirstein, A., Händel, N., Schmid, G. L., Landgraf, K., Berthold, A., Tannert, A., Schaefer, M., Wabitsch, M., Kiess, W., Körner, A., and Garten, A. (2020) A new human adipocyte model with PTEN haploinsufficiency. Adipocyte 9, 290–301 10.1080/21623945.2020.1785083 PMID 32579864

14. Saxton, R. A., and Sabatini, D. M. (2017) mTOR Signaling in Growth, Metabolism, and Disease. Cell 168, 960–976 10.1016/j.cell.2017.02.004 PMID 28283069

15. Simpson, L., and Parsons, R. (2001) PTEN: Life as a tumor suppressor. Experimental cell research 264, 29–41 10.1006/excr.2000.5130 PMID 11237521

16. Schmid, G. L., Kässner, F., Uhlig, H. H., Körner, A., Kratzsch, J., Händel, N., Zepp, F.-P., Kowalzik, F., Laner, A., Starke, S., Wilhelm, F. K., Schuster, S., Viehweger, A., Hirsch, W., Kiess, W., and Garten, A. (2014) Sirolimus treatment of severe PTEN hamartoma tumor syndrome: Case report and in vitro studies. Pediatric research 75, 527–534 10.1038/pr.2013.246 PMID 24366516

17. Huang, W., Queen, N. J., McMurphy, T. B., Ali, S., and Cao, L. (2019) Adipose PTEN regulates adult adipose tissue homeostasis and redistribution via a PTEN-leptin-sympathetic loop. Molecular metabolism 30, 48–60 10.1016/j.molmet.2019.09.008 PMID 31767180

18. Sanchez-Gurmaches, J., Hung, C.-M., Sparks, C. A., Tang, Y., Li, H., and Guertin, D. A. (2012) PTEN loss in the Myf5 lineage redistributes body fat and reveals subsets of white adipocytes that arise from Myf5 precursors. Cell metabolism 16, 348–362 10.1016/j.cmet.2012.08.003 PMID 22940198

19. Filtz, E. A., Emery, A., Lu, H., Forster, C. L., Karasch, C., and Hallstrom, T. C. (2015) Rb1 and Pten Co-Deletion in Osteoblast Precursor Cells Causes Rapid Lipoma Formation in Mice. PLOS ONE 10, e0136729 10.1371/journal.pone.0136729 PMID 26317218

20. Hsieh, S.-C., Chen, N.-T., and Lo, S. H. (2009) Conditional loss of PTEN leads to skeletal abnormalities and lipoma formation. Molecular carcinogenesis 48, 545–552 10.1002/mc.20491 PMID 18973188

21. Chu, C.-Y., Chen, C.-F., Rajendran, R. S., Shen, C.-N., Chen, T.-H., Yen, C.-C., Chuang, C.-K., Lin, D.-S., and Hsiao, C.-D. (2012) Overexpression of Akt1 Enhances Adipogenesis and Leads to Lipoma Formation in Zebrafish. PLOS ONE 7, e36474 10.1371/journal.pone.0036474

22. Fagerberg, L., Hallström, B. M., Oksvold, P., Kampf, C., Djureinovic, D., Odeberg, J., Habuka, M., Tahmasebpoor, S., Danielsson, A., Edlund, K., Asplund, A., Sjöstedt, E., Lundberg, E., Szigyarto, C. A.-K., Skogs, M., Takanen, J. O., Berling, H., Tegel, H., Mulder, J., Nilsson, P., Schwenk, J. M., Lindskog, C., Danielsson, F., Mardinoglu, A., Sivertsson, A., Feilitzen, K. von, Forsberg, M., Zwahlen, M., Olsson, I., Navani, S., Huss, M., Nielsen, J., Ponten, F., and Uhlén, M. (2014) Analysis of the human tissue-specific expression by genome-wide integration of transcriptomics and antibody-based proteomics. Molecular & cellular proteomics: MCP 13, 397–406 10.1074/mcp.M113.035600 PMID 24309898

23. Klingelhutz, A. J., Gourronc, F. A., Chaly, A., Wadkins, D. A., Burand, A. J., Markan, K. R., Idiga, S. O., Wu, M., Potthoff, M. J., and Ankrum, J. A. (2018) Scaffold-free generation of uniform adipose spheroids for metabolism research and drug discovery. Scientific Reports 8, 523 10.1038/s41598-017-19024-z

24. Yoshino, J., Mills, K. F., Yoon, M. J., and Imai, S.-i. (2011) Nicotinamide mononucleotide, a key NAD(+) intermediate, treats the pathophysiology of diet- and age-induced diabetes in mice. Cell metabolism 14, 528–536 10.1016/j.cmet.2011.08.014 PMID 21982712

25. Szklarczyk, D., Gable, A. L., Lyon, D., Junge, A., Wyder, S., Huerta-Cepas, J., Simonovic, M., Doncheva, N. T., Morris, J. H., Bork, P., Jensen, L. J., and Mering, C. von (2019) STRING v11: protein-protein association networks with increased coverage, supporting functional discovery in genome-wide experimental datasets. Nucleic acids research 47, D607–D613 10.1093/nar/gky1131 PMID 30476243

26. Le Duc, D., Lin, C.-C., Popkova, Y., Yang, Z., Akhil, V., Çakir, M. V., Grunewald, S., Simon, J.-C., Dietz, A., Dannenberger, D., Garten, A., Lemke, J. R., Schiller, J., Blüher, M., Nono Nankam, P. A., Rolle-Kampczyk, U., Bergen, M. von, Kelso, J., and Schöneberg, T. (2020) Reduced lipolysis in lipoma phenocopies lipid accumulation in obesity. International journal of obesity (2005) 10.1038/s41366-020-00716-y PMID 33235355

27. Breitfeld, J., Kehr, S., Müller, L., Stadler, P. F., Böttcher, Y., Blüher, M., Stumvoll, M., and Kovacs, P. (2020) Developmentally Driven Changes in Adipogenesis in Different Fat Depots Are Related to Obesity. Frontiers in Endocrinology 11, 138 10.3389/fendo.2020.00138 PMID 32273869

28. Nakae, J., Kitamura, T., Kitamura, Y., Biggs, W. H., Arden, K. C., and Accili, D. (2003) The Forkhead Transcription Factor Foxo1 Regulates Adipocyte Differentiation. Developmental cell 4, 119–129 10.1016/s1534-5807(02)00401-x PMID 12530968

29. Hirsch, J., and Batchelor, B. (1976) Adipose tissue cellularity in human obesity. Clinics in Endocrinology and Metabolism 5, 299–311 10.1016/S0300-595X(76)80023-0

30. García, Z., Kumar, A., Marqués, M., Cortés, I., and Carrera, A. C. (2006) Phosphoinositide 3-kinase controls early and late events in mammalian cell division. The EMBO journal 25, 655–661 10.1038/sj.emboj.7600967 PMID 16437156

31. Pal, A., Barber, T. M., van de Bunt, M., Rudge, S. A., Zhang, Q., Lachlan, K. L., Cooper, N. S., Linden, H., Levy, J. C., Wakelam, M. J. O., Walker, L., Karpe, F., and Gloyn, A. L. (2012) PTEN mutations as a cause of constitutive insulin sensitivity and obesity. The New England journal of medicine 367, 1002–1011 10.1056/NEJMoa1113966 PMID 22970944

32. Tang, E. D., Nuñez, G., Barr, F. G., and Guan, K. L. (1999) Negative regulation of the forkhead transcription factor FKHR by Akt. The Journal of biological chemistry 274, 16741–16746 10.1074/jbc.274.24.16741 PMID 10358014

33. Deng, X., Zhang, W., O-Sullivan, I., Williams, J. B., Dong, Q., Park, E. A., Raghow, R., Unterman, T. G., and Elam, M. B. (2012) FoxO1 inhibits sterol regulatory element-binding protein-1c (SREBP-1c) gene expression via transcription factors Sp1 and SREBP-1c. The Journal of biological chemistry 287, 20132–20143 10.1074/jbc.M112.347211 PMID 22511764

34. Schaum, N., Lehallier, B., Hahn, O., Pálovics, R., Hosseinzadeh, S., Lee, S. E., Sit, R., Lee, D. P., Losada, P. M., Zardeneta, M. E., Fehlmann, T., Webber, J. T., McGeever, A., Calcuttawala, K., Zhang, H., Berdnik, D., Mathur, V., Tan, W., Zee, A., Tan, M., Pisco, A. O., Karkanias, J., Neff, N. F., Keller, A., Darmanis, S., Quake, S. R., and Wyss-Coray, T. (2020) Ageing hallmarks exhibit organ-specific temporal signatures. Nature 153, 1194 10.1038/s41586-020-2499-y

35. Choudhery, M. S., Badowski, M., Muise, A., Pierce, J., and Harris, D. T. (2014) Donor age negatively impacts adipose tissue-derived mesenchymal stem cell expansion and differentiation. Journal of translational medicine 12, 8 10.1186/1479-5876-12-8 PMID 24397850

36. Ramaswamy, S., Nakamura, N., Vazquez, F., Batt, D. B., Perera, S., Roberts, T. M., and Sellers, W. R. (1999) Regulation of G1 progression by the PTEN tumor suppressor protein is linked to inhibition of the phosphatidylinositol 3-kinase/Akt pathway. PNAS 96, 2110–2115 10.1073/pnas.96.5.2110

37. Tu, Q., Hao, J., Zhou, X., Yan, L., Dai, H., Sun, B., Yang, D., An, S., Lv, L., Jiao, B., Chen, C., Lai, R., Shi, P., and Zhao, X. (2018) CDKN2B deletion is essential for pancreatic cancer development instead of unmeaningful co-deletion due to juxtaposition to CDKN2A. Oncogene 37, 128–138 10.1038/onc.2017.316 PMID 28892048

38. Feng, Y., Zhou, L., Sun, X., and Li, Q. (2017) Homeodomain-interacting protein kinase 2 (HIPK2): a promising target for anti-cancer therapies. Oncotarget 8, 20452–20461 10.18632/oncotarget.14723 PMID 28107201

39. Igarashi, M., Miura, M., Williams, E., Jaksch, F., Kadowaki, T., Yamauchi, T., and Guarente, L. (2019) NAD+ supplementation rejuvenates aged gut adult stem cells. Aging cell 18, e12935 10.1111/acel.12935 PMID 30917412

40. Berniakovich, I., Trinei, M., Stendardo, M., Migliaccio, E., Minucci, S., Bernardi, P., Pelicci, P. G., and Giorgio, M. (2008) p66Shc-generated oxidative signal promotes fat accumulation. The Journal of biological chemistry 283, 34283–34293 10.1074/jbc.M804362200 PMID 18838380

41. Eckel-Mahan, K., Ribas Latre, A., and Kolonin, M. G. (2020) Adipose Stromal Cell Expansion and Exhaustion: Mechanisms and Consequences. Cells 9 10.3390/cells9040863 PMID 32252348

42. Schumacher, B., van der Pluijm, I., Moorhouse, M. J., Kosteas, T., Robinson, A. R., Suh, Y., Breit, T. M., van Steeg, H., Niedernhofer, L. J., van Ijcken, W., Bartke, A., Spindler, S. R., Hoeijmakers, J. H. J., van der Horst, G. T. J., and Garinis, G. A. (2008) Delayed and accelerated aging share common longevity assurance mechanisms. PLoS genetics 4, e1000161 10.1371/journal.pgen.1000161 PMID 18704162

43. Kässner, F., Sauer, T., Penke, M., Richter, S., Landgraf, K., Körner, A., Kiess, W., Händel, N., and Garten, A. (2018) Simvastatin induces apoptosis in PTEN-haploinsufficient lipoma cells. International journal of molecular medicine 41, 3691–3698 10.3892/ijmm.2018.3568 PMID 29568880

44. Estève, D., Boulet, N., Volat, F., Zakaroff-Girard, A., Ledoux, S., Coupaye, M., Decaunes, P., Belles, C., Gaits-Iacovoni, F., Iacovoni, J. S., Rémaury, A., Castel, B., Ferrara, P., Heymes, C., Lafontan, M., Bouloumié, A., and Galitzky, J. (2015) Human white and brite adipogenesis is supported by MSCA1 and is impaired by immune cells. Stem cells (Dayton, Ohio) 33, 1277–1291 10.1002/stem.1916 PMID 25523907

45. Fischer-Posovszky, P., Newell, F. S., Wabitsch, M., and Tornqvist, H. E. (2008) Human SGBS cells - a unique tool for studies of human fat cell biology. Obesity facts 1, 184–189 10.1159/000145784 PMID 20054179

46. Schneider, C. A., Rasband, W. S., and Eliceiri, K. W. (2012) NIH Image to ImageJ: 25 years of Image Analysis. Nature methods 9, 671–675 PMID 22930834

47. Sanjabi, S., Williams, K. J., Saccani, S., Zhou, L., Hoffmann, A., Ghosh, G., Gerondakis, S., Natoli, G., and Smale, S. T. (2005) A c-Rel subdomain responsible for enhanced DNA-binding affinity and selective gene activation. Genes & development 19, 2138–2151 10.1101/gad.1329805 PMID 16166378

48. (27.11.2020) UCSC Genome Browser Home. https://genome.ucsc.edu/index.html

49. Afgan, E., Baker, D., Batut, B., van den Beek, M., Bouvier, D., Cech, M., Chilton, J., Clements, D., Coraor, N., Grüning, B. A., Guerler, A., Hillman-Jackson, J., Hiltemann, S., Jalili, V., Rasche, H., Soranzo, N., Goecks, J., Taylor, J., Nekrutenko, A., and Blankenberg, D. (2018) The Galaxy platform for accessible, reproducible and collaborative biomedical analyses: 2018 update. Nucleic acids research 46, W537–W544 10.1093/nar/gky379 PMID 29790989

50. Andrews, S. (2010) Babraham Bioinformatics - FastQC A Quality Control tool for High Throughput Sequence Data. https://www.bioinformatics.babraham.ac.uk/projects/fastqc/

51. Krueger, F. (2012) Babraham Bioinformatics - Trim Galore! http://www.bioinformatics.babraham.ac.uk/projects/trim_galore/

52. Hoffmann, S., Otto, C., Kurtz, S., Sharma, C. M., Khaitovich, P., Vogel, J., Stadler, P. F., and Hackermüller, J. (2009) Fast mapping of short sequences with mismatches, insertions and deletions using index structures. PLoS computational biology 5, e1000502 10.1371/journal.pcbi.1000502 PMID 19750212

53. Love, M. I., Huber, W., and Anders, S. (2014) Moderated estimation of fold change and dispersion for RNA-seq data with DESeq2. Genome biology 15, 550 10.1186/s13059-014-0550-8 PMID 25516281

54. Stephens, M. (2017) False discovery rates: a new deal. Biostatistics (Oxford, England) 18, 275–294 10.1093/biostatistics/kxw041 PMID 27756721

55. Luo, W., and Brouwer, C. (2013) Pathview: an R/Bioconductor package for pathway-based data integration and visualization. Bioinformatics (Oxford, England) 29, 1830–1831 10.1093/bioinformatics/btt285 PMID 23740750

56. Tsai, C.-A., Chen, Y.-J., and Chen, J. J. (2003) Testing for differentially expressed genes with microarray data. Nucleic acids research 31, e52 PMID 12711697

