## Supplementary material for "PTEN regulates adipocyte progenitor growth, differentiation and replicative aging": Figure S4

### PTEN KD

#### PTEN/α-Tubulin

**Gel 1**

| Lane | cell strain | passage | generation | siRNA | α-Tubulin | PTEN | PTEN/Tubulin | Fold |
|---|---|---|---|---|---|---|---|---|
| 1 | SVF#10 | 15 | 17.5 | control | 29685.8 | 30858.8 | 1.0395 | 1 |
| 2 | SVF#10 | 15 | 17.5 | PTEN | 25892.3 | 6868.3 | 0.2653 | 0.255 |
| 3 | SVF#10 | 19 | 20.5 | control | 26662.7 | 25792.9 | 0.9674 | 1 |
| 4 | SVF#10 | 19 | 20.5 | PTEN | 25048.1 | 14121.5 | 0.5638 | 0.583 |
| 5 | SVF#10 | 21 | 21.5 | control | 21221.5 | 16851.7 | 0.7941 | 1 |
| 6 | SVF#10 | 21 | 21.5 | PTEN | 24097.4 | 1199.8 | 0.0498 | 0.063 |

PTEN

α-Tubulin

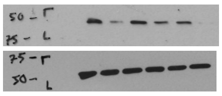

**Gel 2**

| Lane | cell strain | passage | generation | siRNA | α-Tubulin | PTEN | PTEN/Tubulin | Fold |
|---|---|---|---|---|---|---|---|---|
| 1 | SVF#6 | 20 | 19.5 | control | 28432.6 | 34222.4 | 1.2036 | 1 |
| 2 | SVF#6 | 20 | 19.5 | PTEN | 28163.5 | 13651.6 | 0.4847 | 0.403 |
| 3 | SVF#6 | 23 | 25 | control | 28809.4 | 23717.6 | 0.8233 | 1 |
| 4 | SVF#6 | 23 | 25 | PTEN | 26735.5 | 13076.3 | 0.4891 | 0.594 |
| 5 | SVF#6 | 13 | 15 | control | 32007.1 | 36336.6 | 1.1353 | 1 |
| 6 | SVF#6 | 13 | 15 | PTEN | 27552.4 | 13760.5 | 0.4994 | 0.44 |

PTEN

α-Tubulin

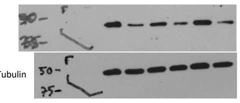

**Gel 3**

| Lane | cell strain | passage | generation | siRNA | α-Tubulin | PTEN | PTEN/Tubulin | Fold |
|---|---|---|---|---|---|---|---|---|
| 1 | SVF#9 | 12 | 14.5 | control | 26326.2 | 34254.7 | 1.3012 | 1 |
| 2 | SVF#9 | 12 | 14.5 | PTEN | 23683.8 | 18228.0 | 0.7696 | 0.592 |
| 3 | SVF#9 | 16 | 19.5 | control | 26977.9 | 38985.1 | 1.4451 | 1 |
| 4 | SVF#9 | 16 | 19.5 | PTEN | 23271.4 | 4514.9 | 0.1940 | 0.134 |
| 5 | SVF#9 | 19 | 23 | control | 22783.1 | 25602.5 | 1.1237 | 1 |
| 6 | SVF#9 | 19 | 23 | PTEN | 25241.9 | 6661.7 | 0.2639 | 0.235 |

PTEN

α-Tubulin

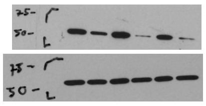

**Gel 4**

| Lane | cell strain | passage | generation | siRNA | α-Tubulin | PTEN | PTEN/Tubulin | Fold |
|---|---|---|---|---|---|---|---|---|
| 1 | SVF#10 | 18 | 27 | control | 25150.6 | 20024.3 | 0.7962 | 1 |
| 2 | SVF#10 | 18 | 27 | PTEN | 29801.7 | 17714.1 | 0.5944 | 0.747 |
| 3 | SVF#9 | 25 | 29.5 | control | 28390.2 | 24607.5 | 0.8668 | 1 |
| 4 | SVF#9 | 25 | 29.5 | PTEN | 31141.2 | 14283.6 | 0.4587 | 0.529 |
| 5 | SVF#10 | 16 | 24.5 | control | 42170.0 | 53220.4 | 1.2620 | 1 |
| 6 | SVF#10 | 16 | 24.5 | PTEN | 49634.0 | 41527.7 | 0.8367 | 0.663 |
| 7 | SVF#10 | 19 | 28 | control | 36816.0 | 36749.9 | 0.9982 | 1 |
| 8 | SVF#10 | 19 | 28 | PTEN | 33632.1 | 18851.5 | 0.5605 | 0.562 |

PTEN

α-Tubulin

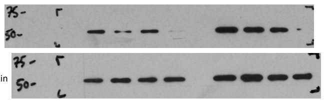

**Gel 5**

| Lane | cell strain | passage | generation | siRNA | α-Tubulin | PTEN | PTEN/Tubulin | Fold |
|---|---|---|---|---|---|---|---|---|
| 1 | SVF#10 | 22 | 31.5 | control | 25567.1 | 26008.5 | 1.0173 | 1 |
| 2 | SVF#10 | 22 | 31.5 | PTEN | 22026.5 | 14362.9 | 0.6521 | 0.641 |
| 3 | SVF#8 | 6 | 11.5 | control | 23997.8 | 27608.4 | 1.1505 | 1 |
| 4 | SVF#8 | 6 | 11.5 | PTEN | 22733.3 | 20224.4 | 0.8896 | 0.773 |
| 5 | SVF#6 | 19 | 27.5 | control | 25526.2 | 28333.1 | 1.1100 | 1 |
| 6 | SVF#6 | 19 | 27.5 | PTEN | 22336.8 | 14500.3 | 0.6492 | 0.585 |
| 7 | SVF#6 | 10 | 16.5 | control | 32681.5 | 48733.2 | 1.4912 | 1 |
| 8 | SVF#6 | 10 | 16.5 | PTEN | 31490.7 | 31924.4 | 1.0138 | 0.68 |
| 9 | SVF#6 | 14 | 21 | control | 23326.8 | 21936.5 | 0.9404 | 1 |
| 10 | SVF#6 | 14 | 21 | PTEN | 27925.7 | 10446.5 | 0.3741 | 0.398 |

PTEN

α-Tubulin

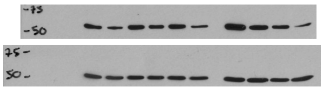

#### pAKT/AKT

**Gel 1**

| Lane | cell strain | passage | generation | siRNA | pAKT | AKT | pAKT/AKT | Fold |
|---|---|---|---|---|---|---|---|---|
| 1 | SVF#10 | 22 | 31.5 | control | 8394.953 | 24410.3 | 0.3439 | 1 |
| 2 | SVF#10 | 22 | 31.5 | PTEN | 22735.38 | 28621.7 | 0.7943 | 2.31 |
| 3 | SVF#8 | 6 | 11.5 | control | 412.92 | 27179.5 | 0.0152 | 1 |
| 4 | SVF#8 | 6 | 11.5 | PTEN | 915.749 | 26966.7 | 0.0340 | 2.235 |
| 5 | SVF#6 | 19 | 27.5 | control | 136.95 | 27450.6 | 0.0050 | 1 |
| 6 | SVF#6 | 19 | 27.5 | PTEN | 433.263 | 28802 | 0.0150 | 3.015 |
| 7 | SVF#6 | 10 | 16.5 | control | 896.477 | 45455.4 | 0.0197 | 1 |
| 8 | SVF#6 | 10 | 16.5 | PTEN | 37526.342 | 50902.8 | 0.7372 | 37.38 |
| 9 | SVF#6 | 14 | 21 | control | 25207.38 | 40361.9 | 0.6245 | 1 |
| 10 | SVF#6 | 14 | 21 | PTEN | 33553.664 | 34367.8 | 0.9763 | 1.563 |

pAKT (T308)

AKT

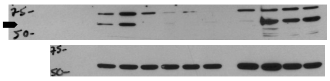

**Gel 2**

| Lane | cell strain | passage | generation | siRNA | AKT | pAKT | pAKT/AKT | Fold |
|---|---|---|---|---|---|---|---|---|
| 1 | SVF#10 | 18 | 27 | control | 25284.451 | | | |
| 2 | SVF#10 | 18 | 27 | PTEN | 31423.43 | | | |
| 3 | SVF#9 | 25 | 29.5 | control | 20953.945 | | | |
| 4 | SVF#9 | 25 | 29.5 | PTEN | 24168.702 | | | |
| 5 | SVF#10 | 16 | 24.5 | control | 36576.794 | 191.192 | 0.0052 | 1 |
| 6 | SVF#10 | 16 | 24.5 | PTEN | 44530.957 | 23790 | 0.5342 | 102.2 |
| 7 | SVF#10 | 19 | 28 | control | 39983.886 | 28464.9 | 0.7119 | 1 |
| 8 | SVF#10 | 19 | 28 | PTEN | 19359.48 | 37363.3 | 1.9300 | 2.711 |

pAKT (T308)

AKT

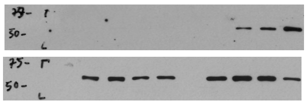

#### pS6/α-Tubulin

**Gel 1**

| Lane | cell strain | passage | generation | siRNA | α-Tubulin | pS6 | pS6/Tubulin | Fold |
|---|---|---|---|---|---|---|---|---|
| 1 | SVF#10 | 15 | 17.5 | control | 29685.765 | 1591.79 | 0.0536 | 1 |
| 2 | SVF#10 | 15 | 17.5 | PTEN | 25892.258 | 14296.5 | 0.5522 | 10.3 |
| 3 | SVF#10 | 19 | 20.5 | control | 26662.673 | 6980.95 | 0.2618 | 1 |
| 4 | SVF#10 | 19 | 20.5 | PTEN | 25048.116 | 26784.3 | 1.0693 | 4.084 |
| 5 | SVF#10 | 21 | 21.5 | control | 21221.459 | 985.891 | 0.0465 | 1 |
| 6 | SVF#10 | 21 | 21.5 | PTEN | 24097.409 | 15372.4 | 0.6379 | 13.73 |

pS6 (Ser235/236)

α-Tubulin

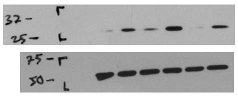

**Gel 2**

| Lane | cell strain | passage | generation | siRNA | α-Tubulin | pS6 | pS6/Tubulin | Fold |
|---|---|---|---|---|---|---|---|---|
| 1 | SVF#10 | 22 | 31.5 | control | 25567.078 | 64222.7 | 2.5119 | 1 |
| 2 | SVF#10 | 22 | 31.5 | PTEN | 22026.501 | 73514.5 | 3.3375 | 1.329 |
| 3 | SVF#8 | 6 | 11.5 | control | 23997.794 | 2306.21 | 0.0961 | 1 |
| 4 | SVF#8 | 6 | 11.5 | PTEN | 22733.258 | 6170.49 | 0.2714 | 2.824 |
| 5 | SVF#6 | 19 | 27.5 | control | 25526.208 | | | |
| 6 | SVF#6 | 19 | 27.5 | PTEN | 22336.773 | | | |
| 7 | SVF#6 | 10 | 16.5 | control | 32681.501 | 1563.01 | 0.0478 | 1 |
| 8 | SVF#6 | 10 | 16.5 | PTEN | 31490.723 | 5657.71 | 0.1797 | 3.757 |
| 9 | SVF#6 | 14 | 21 | control | 23326.752 | 239.192 | 0.0103 | 1 |
| 10 | SVF#6 | 14 | 21 | PTEN | 27925.693 | 15612.9 | 0.5591 | 54.52 |

pS6 (Ser235/236)

α-Tubulin

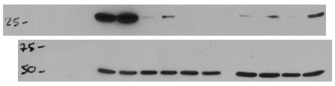

**Gel 3**

| Lane | cell strain | passage | generation | siRNA | α-Tubulin | pS6 | pS6/Tubulin | Fold |
|---|---|---|---|---|---|---|---|---|
| 1 | SVF#10 | 18 | 27 | control | 25150.572 | 471.556 | 0.0187 | 1 |
| 2 | SVF#10 | 18 | 27 | PTEN | 29801.744 | 24020.1 | 0.8060 | 42.99 |
| 3 | SVF#9 | 25 | 29.5 | control | 28390.179 | 421.435 | 0.0148 | 1 |
| 4 | SVF#9 | 25 | 29.5 | PTEN | 31141.229 | 2097.5 | 0.0674 | 4.537 |
| 5 | SVF#10 | 16 | 24.5 | control | 42169.957 | 5913.85 | 0.1402 | 1 |
| 6 | SVF#10 | 16 | 24.5 | PTEN | 49634.049 | 29104.5 | 0.5864 | 4.181 |
| 7 | SVF#10 | 19 | 28 | control | 36816.007 | 41326.6 | 1.1225 | 1 |
| 8 | SVF#10 | 19 | 28 | PTEN | 33632.057 | 40305.2 | 1.1984 | 1.068 |

pS6 (Ser235/236)

α-Tubulin

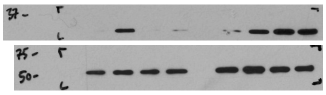

#### pFOXO1/α-Tubulin

| Lane | cell strain | passage | generation | siRNA | α-Tubulin | pFOXO1 | pFOXO1/ Tubulin | Fold |
|---|---|---|---|---|---|---|---|---|
| 1 | SVF#6 | 23 | 25 | control | 23465.723 | 1664.86 | 0.0709 | 1 |
| 2 | SVF#6 | 23 | 25 | PTEN | 25824.551 | 9051.15 | 0.3505 | 4.94 |
| 3 | SVF#9 | 16 | 19.5 | control | 19014.723 | 5445.25 | 0.2864 | 1 |
| 4 | SVF#9 | 16 | 19.5 | PTEN | 21137.915 | 9373.2 | 0.4434 | 1.548 |
| 5 | SVF#10 | 19 | 28 | control | 18774.551 | 219.778 | 0.0117 | 1 |
| 6 | SVF#10 | 19 | 28 | PTEN | 18489.844 | 2879.62 | 0.1557 | 13.3 |

pFOXO1

α-Tubulin

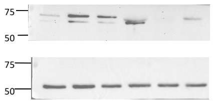


#### SREBP1/α-Tubulin

| Lane | cell strain | passage | generation | siRNA | α-Tubulin | SREBP1 | SREBP1/ Tubulin | Fold |
|---|---|---|---|---|---|---|---|---|
| 1 | SVF#10 | 19 | 20.5 | control | 6903.933 | 9476.12 | 1.3726 | 1 |
| 2 | SVF#10 | 19 | 20.5 | PTEN | 8575.296 | 19465.1 | 2.2699 | 1.654 |
| 3 | SVF#6 | 20 | 19.5 | control | 5463.64 | 6880.17 | 1.2593 | 1 |
| 4 | SVF#6 | 20 | 19.5 | PTEN | 5284.004 | 11169.2 | 2.1138 | 1.679 |
| 5 | SVF#9 | 12 | 14.5 | control | 3411.983 | 6350.1 | 1.8611 | 1 |
| 6 | SVF#9 | 12 | 14.5 | PTEN | 2969.569 | 6134.44 | 2.0658 | 1.11 |
| 7 | SVF#8 | 13 | 22.5 | control | 4734.225 | 3256.18 | 0.6878 | 1 |
| 8 | SVF#8 | 13 | 22.5 | PTEN | 5846.175 | 6014.9 | 1.0289 | 1.496 |

SREBP1

α-Tubulin

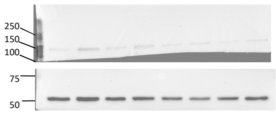
