## Supplementary material for "PTEN regulates adipocyte progenitor growth, differentiation and replicative aging": Figure S5

### PTEN CRISPR SVF

#### PTEN/α-Tubulin

**Gel 1**

| Lane | Cells | guideRNA | α-Tubulin | PTEN | PTEN/Tubulin | Fold |
|---|---|---|---|---|---|---|
| 1 | CRISPR SVF#1 | control | 44983.5 | 35283.2 | 0.7844 | 1 |
| 2 | CRISPR SVF#1 | PTEN #3+4 | 41204.8 | 23047.4 | 0.5593 | 0.7131 |
| 3 | CRISPR SVF#3 | control | 39369.1 | 34844.8 | 0.8851 | 1 |
| 4 | CRISPR SVF#3 | PTEN #3+4 | 32432.8 | 2196.4 | 0.0677 | 0.0765 |

PTEN

α-Tubulin

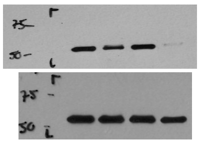

**Gel 2**

| Lane | Cells | guideRNA | α-Tubulin | PTEN | PTEN/Tubulin | Fold |
|---|---|---|---|---|---|---|
| 1 | CRISPR SVF#9 | control | 30284.5 | 37275.7 | 1.2309 | 1 |
| 2 | CRISPR SVF#9 | PTEN #3+4 | 25771.4 | 40277.4 | 1.5629 | 1.2697 |

PTEN

α-Tubulin

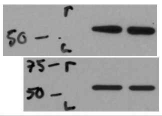

**Gel 3**

| Lane | Cells | guideRNA | GAPDH | PTEN | PTEN/GAPDH | Fold |
|---|---|---|---|---|---|---|
| 1 | CRISPR SVF#6 | control | 46613.2 | 25935.4 | 0.5564 | 1 |
| 2 | CRISPR SVF#6 | control | 37796.5 | 20527.1 | 0.5431 | 0.9761 |
| 3 | CRISPR SVF#6 | PTEN #3 | 48217.2 | 22996.5 | 0.4769 | 0.8572 |
| 4 | CRISPR SVF#6 | PTEN #3+4 | 50484.9 | 16209.0 | 0.3211 | 0.5770 |

PTEN

GAPDH

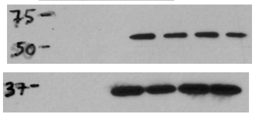

#### pAKT/AKT

**Gel 1**

| Lane | Cells | guideRNA | pAKT | AKT | pAKT/AKT | Fold |
|---|---|---|---|---|---|---|
| 1 | CRISPR SVF#1 | control | 24115.673 | 47316.22 | 0.50967032 | 1 |
| 2 | CRISPR SVF#1 | PTEN #3+4 | 15713.116 | 42532.735 | 0.369435824 | 0.724852536 |
| 3 | CRISPR SVF#3 | control | 16171.53 | 46697.271 | 0.346305676 | 1 |
| 4 | CRISPR SVF#3 | PTEN #3+4 | 28030.016 | 43814.614 | 0.639741252 | 1.847331118 |

pAKT (T308)

AKT

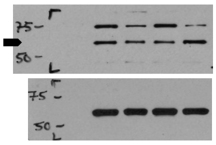

**Gel 2**

| Lane | Cells | guideRNA | pAKT | AKT | pAKT/AKT | Fold |
|---|---|---|---|---|---|---|
| 1 | CRISPR SVF#9 | control | 18924.9 | 45458.3 | 0.4163 | 1 |
| 2 | CRISPR SVF#9 | PTEN #3+4 | 40323.7 | 41352.1 | 0.9751 | 2.3423 |

pAKT (T308)

AKT

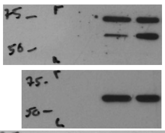

**Gel 3**

| Lane | Cells | guideRNA | pAKT | AKT | pAKT/AKT | Fold |
|---|---|---|---|---|---|---|
| 1 | CRISPR SVF#6 | control | 1557.7 | 34414.0 | 0.0453 | 1 |
| 2 | CRISPR SVF#6 | control | 1137.2 | 24325.5 | 0.0468 | 1.0329 |
| 3 | CRISPR SVF#6 | PTEN #3 | 18959.9 | 32016.1 | 0.5922 | 13.0837 |
| 4 | CRISPR SVF#6 | PTEN #3+4 | 43735.2 | 37969.8 | 1.1518 | 25.4480 |

pAKT (T308)

AKT

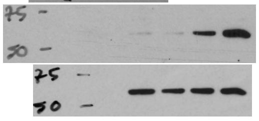

#### pS6/α-Tubulin

**Gel 1**

| Lane | Cells | guideRNA | α-Tubulin | pS6 | pS6/Tubulin | Fold |
|---|---|---|---|---|---|---|
| 1 | CRISPR SVF#1 | control | 19917.5 | 29887.6 | 1.5006 | 1 |
| 2 | CRISPR SVF#1 | PTEN #3+4 | 20513.1 | 47436.4 | 2.3125 | 1.5411 |
| 3 | CRISPR SVF#3 | control | 24033.8 | 20507.5 | 0.8533 | 1 |
| 4 | CRISPR SVF#3 | PTEN #3+4 | 22460.1 | 49393.8 | 2.1992 | 2.5773 |

pS6 (Ser235/236)

α-Tubulin

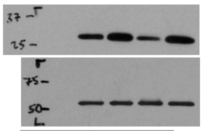

**Gel 2**

| Lane | Cells | guideRNA | α-Tubulin | pS6 | pS6/Tubulin | Fold |
|---|---|---|---|---|---|---|
| 1 | CRISPR SVF#9 | control | 30284.5 | 527.7 | 0.0174 | 1 |
| 2 | CRISPR SVF#9 | PTEN #3+4 | 25771.4 | 19182.4 | 0.7443 | 42.7126 |

pS6 (Ser235/236)

α-Tubulin

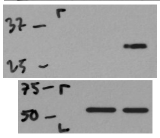

**Gel 3**

| Lane | Cells | guideRNA | GAPDH | pS6 | pS6/GAPDH | Fold |
|---|---|---|---|---|---|---|
| 1 | CRISPR SVF#6 | control | 55297.3 | 8164.4 | 0.1476 | 1 |
| 2 | CRISPR SVF#6 | control | 41454.0 | 3526.6 | 0.0851 | 0.5762 |
| 3 | CRISPR SVF#6 | PTEN #3 | 35198.0 | 23966.1 | 0.6809 | 4.6117 |
| 4 | CRISPR SVF#6 | PTEN #3+4 | 37901.3 | 49345.4 | 1.3019 | 8.8180 |

pS6 (Ser235/236)

GAPDH

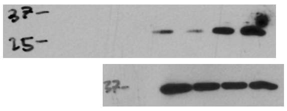

### Constitutive active FOXO1 overexpression in CRISPR SVF

#### FOXO1 active/α-Tubulin

**Gel 1**

| Lane | Cells | guideRNA | Plasmid | α-Tubulin | FOXO1 (FLAG) | FOXO1/Tubulin | Fold |
|---|---|---|---|---|---|---|---|
| 1 | CRISPR SVF#6 p16 | PTEN #3+4 | control | 7418.8 | 476.2 | 0.0642 | 1 |
| 2 | CRISPR SVF#6 p16 | PTEN #3+4 | FOXO1 active | 9221.7 | 13958.1 | 1.5136 | 23.58 |
| 3 | CRISPR SVF#6 p17 | PTEN #3+4 | control | 7703.0 | 246.1 | 0.0320 | 1 |
| 4 | CRISPR SVF#6 p17 | PTEN #3+4 | FOXO1 active | 7416.1 | 7366.7 | 0.9933 | 31.09 |

FOXO1 (FLAG)

α-Tubulin

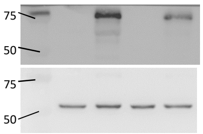


**Gel 2**

| Lane | Cells | guideRNA | Plasmid | α-Tubulin | FOXO1 (FLAG) | FOXO1/Tubulin | Fold |
|---|---|---|---|---|---|---|---|
| 1 | CRISPR SVF#6 p23 | PTEN #3+4 | control | 5765.4 | 19.0 | 0.0033 | 1 |
| 2 | CRISPR SVF#6 p23 | PTEN #3+4 | FOXO1 active | 6649.6 | 9061.5 | 1.3627 | 414.60 |

FOXO1 (FLAG)

α-Tubulin

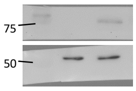


#### SREBP1/α-Tubulin

**Gel 1**

| Lane | Cells | guideRNA | Plasmid | α-Tubulin | SREBP1 | SREBP1/Tubulin | Fold |
|---|---|---|---|---|---|---|---|
| 1 | CRISPR SVF#6 p16 | PTEN #3+4 | control | 7418.8 | 5959.8 | 0.8033 | 1 |
| 2 | CRISPR SVF#6 p16 | PTEN #3+4 | FOXO1 active | 9221.7 | 3821.9 | 0.4145 | 0.52 |
| 3 | CRISPR SVF#6 p17 | PTEN #3+4 | control | 7703.0 | 7921.5 | 1.0284 | 1 |
| 4 | CRISPR SVF#6 p17 | PTEN #3+4 | FOXO1 active | 7416.1 | 5395.6 | 0.7276 | 0.71 |

SREBP1

α-Tubulin

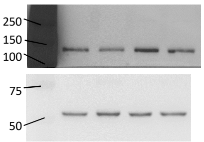


**Gel 2**

| Lane | Cells | guideRNA | Plasmid | α-Tubulin | SREBP1 | SREBP1/Tubulin | Fold |
|---|---|---|---|---|---|---|---|
| 1 | CRISPR SVF#6 p23 | PTEN #3+4 | control | 5765.4 | 3939.5 | 0.6833 | 1 |
| 2 | CRISPR SVF#6 p23 | PTEN #3+4 | FOXO1 active | 6649.6 | 1095.8 | 0.1648 | 0.24 |

SREBP1

α-Tubulin

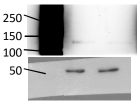
