## Supplementary material for "PTEN regulates adipocyte progenitor growth, differentiation and replicative aging": Figure S6

### PTEN KD

#### NAMPT/α-Tubulin

**Gel 1**

| Lane | cell strain | passage | generation | siRNA | α-Tubulin | NAMPT | NAMPT/Tubulin | Fold |
|---|---|---|---|---|---|---|---|---|
| 1 | SVF#10 | 15 | 17.5 | control | 29685.8 | 11692.5 | 0.3939 | 1 |
| 2 | SVF#10 | 15 | 17.5 | PTEN | 25892.3 | 25617.6 | 0.9894 | 2.5119 |
| 3 | SVF#10 | 19 | 20.5 | control | 26662.7 | 11961.0 | 0.4486 | 1 |
| 4 | SVF#10 | 19 | 20.5 | PTEN | 25048.1 | 24268.8 | 0.9689 | 2.1598 |
| 5 | SVF#10 | 21 | 21.5 | control | 21221.5 | 19353.1 | 0.9120 | 1 |
| 6 | SVF#10 | 21 | 21.5 | PTEN | 24097.4 | 23821.2 | 0.9885 | 1.0840 |

NAMPT

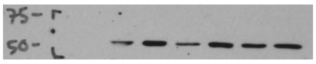

α-Tubulin

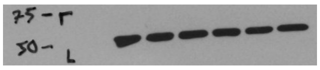

**Gel 2**

| Lane | cell strain | passage | generation | siRNA | α-Tubulin | NAMPT | NAMPT/Tubulin | Fold |
|---|---|---|---|---|---|---|---|---|
| 1 | SVF#10 | 18 | 27 | control | 25150.572 | 18085.338 | 0.7191 | 1 |
| 2 | SVF#10 | 18 | 27 | PTEN | 29801.744 | 29915.401 | 1.0038 | 1.3960 |
| 3 | SVF#9 | 25 | 29.5 | control | 28390.179 | 20071.702 | 0.7070 | 1 |
| 4 | SVF#9 | 25 | 29.5 | PTEN | 31141.229 | 31966.886 | 1.0265 | 1.4519 |
| 5 | SVF#10 | 16 | 24.5 | control | 42169.957 | 33930.229 | 0.8046 | 1 |
| 6 | SVF#10 | 16 | 24.5 | PTEN | 49634.049 | 50054.726 | 1.0085 | 1.2534 |
| 7 | SVF#10 | 19 | 28 | control | 36816.007 | 37188.907 | 1.0101 | 1 |
| 8 | SVF#10 | 19 | 28 | PTEN | 33632.057 | 43514.605 | 1.2938 | 1.2809 |

NAMPT

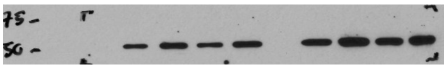

α-Tubulin

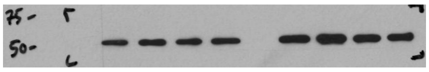

#### p21/α-Tubulin

**Gel 1**

| Lane | cell strain | passage | generation | siRNA | α-Tubulin | p21 | p21/Tubulin | Fold |
|---|---|---|---|---|---|---|---|---|
| 1 | SVF#10 | 18 | 27 | control | 25150.572 | 12232.974 | 0.4864 | 1 |
| 2 | SVF#10 | 18 | 27 | PTEN | 29801.744 | 5668.246 | 0.1902 | 0.3910 |
| 3 | SVF#9 | 25 | 29.5 | control | 28390.179 | 11907.723 | 0.4194 | 1 |
| 4 | SVF#9 | 25 | 29.5 | PTEN | 31141.229 | 8778.782 | 0.2819 | 0.6721 |
| 5 | SVF#10 | 16 | 24.5 | control | 42169.957 | 32319.451 | 0.7664 | 1 |
| 6 | SVF#10 | 16 | 24.5 | PTEN | 49634.049 | 15655.045 | 0.3154 | 0.4115 |
| 7 | SVF#10 | 19 | 28 | control | 36816.007 | 25134.057 | 0.6827 | 1 |
| 8 | SVF#10 | 19 | 28 | PTEN | 33632.057 | 4063.104 | 0.1208 | 0.1770 |

p21

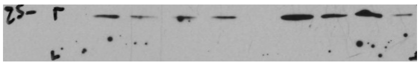

α-Tubulin

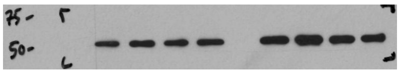

**Gel 2**

| Lane | cell strain | passage | generation | siRNA | α-Tubulin | p21 | p21/Tubulin | Fold |
|---|---|---|---|---|---|---|---|---|
| 1 | SVF#10 | 22 | 31.5 | control | 25002.208 | 16890.459 | 0.6756 | 1 |
| 2 | SVF#10 | 22 | 31.5 | PTEN | 28004.329 | 7717.246 | 0.2756 | 0.4079 |
| 3 | SVF#8 | 6 | 11.5 | control | 25806.501 | 17323.752 | 0.6713 | 1 |
| 4 | SVF#8 | 6 | 11.5 | PTEN | 30205.886 | 7867.246 | 0.2605 | 0.3880 |
| 5 | SVF#6 | 19 | 27.5 | control | 23217.016 | 15644.974 | 0.6739 | 1 |
| 6 | SVF#6 | 19 | 27.5 | PTEN | 24411.501 | 2608.912 | 0.1069 | 0.1586 |
| 7 | SVF#6 | 10 | 16.5 | control | 28875.522 | 4094.447 | 0.1418 | 1 |
| 8 | SVF#6 | 10 | 16.5 | PTEN | 41495.463 | 97.778 | 0.0024 | 0.0166 |

p21

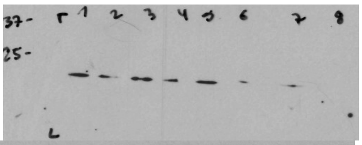

α-Tubulin

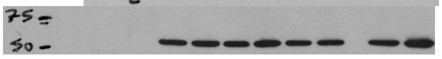

### Longterm culture

**Gel 1**

| Lane | cell strain | passage | generation | days | α-Tubulin | PTEN | PTEN/Tubulin | NAMPT | NAMPT/Tubulin | pAkt | Akt | pAKT/AKT |
|---|---|---|---|---|---|---|---|---|---|---|---|---|
| 1 | SVF#6 | 4 | 5.5 | 11 | 21897.7 | 2176.0 | 0.0994 | 10473.0 | 0.4783 | 56901.7 | 1123.8 | 50.6324 |
| 2 | SVF#6 | 8 | 12 | 24 | 36210.7 | 8694.3 | 0.2401 | 4683.2 | 0.1293 | 42782.3 | 28394.1 | 1.5067 |
| 3 | SVF#6 | 5 | 8.5 | 17 | 34287.5 | 2348.0 | 0.0685 | 5311.5 | 0.1549 | 38720.3 | 27248.1 | 1.4210 |
| 4 | SVF#6 | 12 | 17.5 | 35 | 44295.3 | 20148.3 | 0.4549 | 8349.9 | 0.1885 | 7610.3 | 41283.3 | 0.1843 |
| 5 | SVF#6 | 14 | 19.5 | 39 | 45090.0 | 40007.0 | 0.8873 | 1639.7 | 0.0364 | 16317.2 | 38784.3 | 0.4207 |
| 6 | SVF#6 | 16 | 21.5 | 43 | 36260.4 | 29828.0 | 0.8226 | 544.0 | 0.0150 | 26400.0 | 33258.7 | 0.7938 |
| 7 | SVF#6 | 18 | 22.5 | 45 | 22367.2 | 17480.7 | 0.7815 | 408.3 | 0.0183 | 16106.4 | 15828.7 | 1.0175 |
| 8 | SVF#6 | 19 | 23 | 46 | 31162.5 | 22429.2 | 0.7198 | 3111.9 | 0.0999 | 18106.7 | 34120.4 | 0.5307 |
| 9 | SVF#6 | 21 | 26.5 | 53 | 38233.7 | 23050.0 | 0.6029 | 2255.5 | 0.0590 | 5655.3 | 32125.4 | 0.1760 |
| 10 | SVF#6 | 21 | 27 | 54 | 36219.0 | 18207.8 | 0.5027 | 968.6 | 0.0267 | 4665.3 | 36954.7 | 0.1262 |
| 11 | SVF#6 | 22 | 28.5 | 57 | 36973.0 | 27925.5 | 0.7553 | 1786.5 | 0.0483 | 404.3 | 45943.9 | 0.0088 |
| 12 | SVF#6 | 20 | 40.5 | 81 | 33873.7 | 17538.0 | 0.5177 | 391.0 | 0.0115 | 16532.7 | 17372.9 | 0.9516 |

PTEN

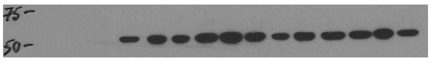

pAKT

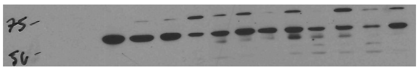

NAMPT

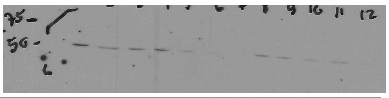

AKT

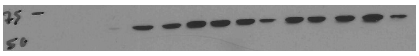

α-Tubulin

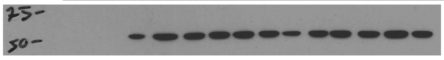

**Gel 2**

| Lane | cell strain | passage | generation | ys in cult | α-Tubulin | PTEN | PTEN/Tubulin | NAMPT | NAMPT/Tubulin | pAkt | Akt | pAKT/AKT |
|---|---|---|---|---|---|---|---|---|---|---|---|---|
| 1 | SVF#8 | 2 | 2 | 4 | 14337.7 | 12774.77 | 0.890991582 | 14437 | 1.006936259 | 29543.81 | 3388.518 | 8.71879978 |
| 2 | SVF#8 | 3 | 3.5 | 7 | 24164.33 | 52807.38 | 2.185344266 | 6341.5 | 0.262433885 | 41677.32 | 16287.34 | 2.55887825 |
| 3 | SVF#8 | 5 | 8.5 | 17 | 42892.41 | 37488.32 | 0.874008245 | 558.92 | 0.013030744 | 56173.32 | 2402.326 | 23.3828881 |
| 4 | SVF#8 | 11 | 18.5 | 37 | 84696.81 | 49400.24 | 0.583259747 | 198.95 | 0.002348967 | 13984.12 | 20216.09 | 0.69173218 |
| 5 | SVF#8 | 12 | 21 | 42 | 69102.91 | 23498.9 | 0.340056591 | 777.65 | 0.011253477 | 42.536 | 14095.02 | 0.0030178 |
| 6 | SVF#8 | 13 | 22.5 | 45 | 57271.12 | 38048.1 | 0.664350549 | 277.9 | 0.004852341 | 8259.246 | 11969.27 | 0.69003757 |
| 7 | SVF#8 | 14 | 25.5 | 51 | 50289.697 | 68391.49 | 1.359950329 | 717.75 | 0.014272287 | 7757.368 | 32665.81 | 0.23747668 |

PTEN

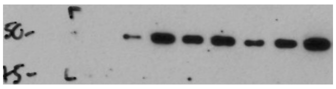

pAKT

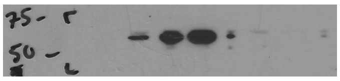

NAMPT

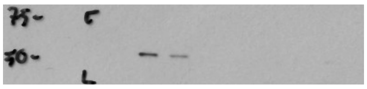

AKT

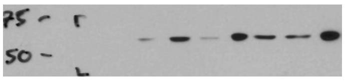

α-Tubulin

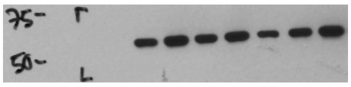

**Gel 3**

| Lane | cell strain | passage | generation | ys in cult | α-Tubulin | PTEN | PTEN/Tubulin | NAMPT | NAMPT/Tubulin | pAkt | Akt | pAKT/AKT |
|---|---|---|---|---|---|---|---|---|---|---|---|---|
| 1 | SVF#9 | 1 | 1 | 2 | 14337.7 | 76.778 | 0.005354973 | 61101 | 4.261530789 | 7211.966 | 2503.69 | 2.88053473 |
| 2 | SVF#9 | 1 | 1.5 | 3 | 24164.33 | 14126.92 | 0.584618733 | 90193 | 3.732468063 | 17232.77 | 13080.48 | 1.31744171 |
| 3 | SVF#9 | 6 | 8.5 | 17 | 42892.41 | 39505.61 | 0.921039643 | 47385 | 1.104740909 | 77.607 | 38965.78 | 0.00199167 |
| 4 | SVF#9 | 9 | 13.5 | 27 | 84696.81 | 65594.47 | 0.774462108 | 32540 | 0.384194281 | 137.728 | 69093.95 | 0.00199334 |
| 5 | SVF#9 | 12 | 14.5 | 29 | 69102.91 | 62116.41 | 0.898897167 | 18288 | 0.264652096 | 29267.32 | 64161.54 | 0.45615052 |
| 6 | SVF#9 | 13 | 16.5 | 33 | 57271.12 | 20776.06 | 0.362766784 | 10323 | 0.180255598 | 9554.147 | 42402.8 | 0.22531878 |
| 7 | SVF#9 | 18 | 24 | 48 | 44063.59 | 25981.66 | 0.589640109 | 18145 | 0.411791005 | 73.192 | 48147.8 | 0.00152015 |

PTEN

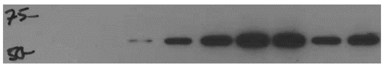

pAKT

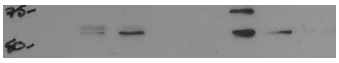

NAMPT

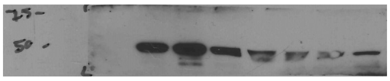

AKT

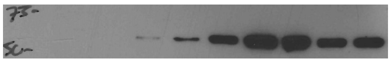

α-Tubulin

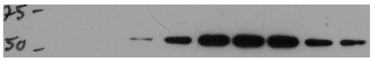

**Gel 4**

| Lane | cell strain | passage | generation | ys in cult | Tubulin | PTEN | PTEN/Tubulin | NAMPT | NAMPT/Tubulin | pAkt | Akt | pAKT/AKT |
|---|---|---|---|---|---|---|---|---|---|---|---|---|
| 1 | SVF#10 | 1 | 3 | 6 | 22996.19 | 9064.288 | 0.394164773 | 54166 | 2.355448446 | 267.021 | 36170.77 | 0.00738223 |
| 2 | SVF#10 | 2 | 4.5 | 9 | 25914.02 | 20824.28 | 0.803591261 | 15888 | 0.613089748 | 29203.35 | 38533.37 | 0.75787168 |
| 3 | SVF#10 | 3 | 5.5 | 11 | 29100.14 | 24330.42 | 0.836092885 | 7606.6 | 0.261394997 | 12482.07 | 44235.22 | 0.28217493 |
| 4 | SVF#10 | 6 | 10 | 20 | 21255.58 | 29670.3 | 1.395882869 | 14390 | 0.677006697 | 36794.81 | 37088.32 | 0.99208619 |
| 5 | SVF#10 | 9 | 14.5 | 29 | 40137.3 | 66097.88 | 1.646794378 | 2711.4 | 0.067553049 | 485.435 | 72107.16 | 0.00673213 |

PTEN

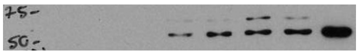

pAKT

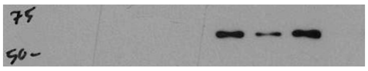

NAMPT

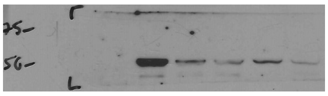

AKT

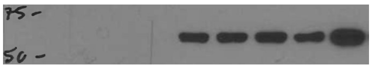

α-Tubulin

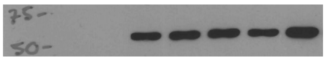
