## Supplementary material for "PTEN regulates adipocyte progenitor growth, differentiation and replicative aging": Table S4

**Table S4.** SVF cell cultures.

| Name | Sex | Age at resection | Origin |
|---|---|---|---|
| SVF#1 | female | 31 | visceral |
| SVF#3 | female | 30 | visceral |
| SVF#5 | female | 33 | visceral |
| SVF#6 | male | 27 | visceral |
| SVF#8 | male | 37 | visceral |
| SVF#9 | female | 32 | visceral |
| SVF#10 | female | 29 | visceral |
| SVF#11S | male | 18 | subcutaneous |
| SVF#12S | male | 24 | subcutaneous |
| SVF#14S | female | 28 | subcutaneous |
