## Supplementary material for "PTEN regulates adipocyte progenitor growth, differentiation and replicative aging": Table S5

**Table S5.** Antibodies used for flow cytometry.

| Marker | Fluorophore | Distributor | Clone | Cat. no |
|---|---|---|---|---|
| MSCA1 | PE | Miltenyi | W8B2 | 130-093-587 |
| CD34 | FITC | Miltenyi | AC136 | 130-113-178 |
| CD14 | PeVio770 | Miltenyi | Tük4 | 130-113-149 |
| CD271 | APC | Miltenyi | ME20.4-1-H4 | 130-113-418 |
| CD8 | PerCP | BD Horizon | SK1 | 345774 |
| CD31 | V450 | BD Horizon | WM59 | 561653 |
| CD45 | BV510 | BioLegend | HI30 | 304036 |
