## Supplementary material for "PTEN regulates adipocyte progenitor growth, differentiation and replicative aging": Table S6

| cells | passage | generation | single cells | viable cells | CD8+ | CD14+ | CD31+ | CD34+ | CD45+ | CD271+ | MSCA1+ |
|---|---|---|---|---|---|---|---|---|---|---|---|
| LipPD1 | 6 | 8 | 97% | 77.60% | none | none | none | none | none | 4.70% | 4.74% |
| LipPD1 | 28 | 34 | 93.60% | 66.70% | none | 6% | none | 3.62% | none | 3.86% | 1.51% |
| SVF#1 | 6 | 19 | 95.30% | 57.00% | none | 2.56% | none | 18.60% | none | 3.67% | none |
| SVF#1 | 12 | 17.5 | 93.70% | 51.90% | none | 2.76% | none | 22.00% | none | 1.58% | 3.00% |
| SVF#1 | 22 | 35.5 | 99.10% | 71.20% | none | none | none | 29.00% | none | none | 1.63% |
| SVF#5 | 0 | 0 | 98.90% | 23.00% | none | 9.10% | 3.04% | 58.70% | 6.18% | 15.80% | 5.96% |
| SVF#5 | 11 | 25.5 | 96.50% | 79.90% | none | none | none | 2% | none | none | none |
| SVF#6 | 0 | 1.5 | 99.60% | 8.04% | none | 2.98% | 2% | none | none | 19% | 19% |
| SVF#6 | 16 | 19 | 96.20% | 79.30% | none | none | none | 12.90% | none | none | 4.46% |
| SVF#6 | 23 | 31 | 97.10% | 74.80% | none | none | none | 38.60% | none | none | 2.44% |
| SVF#8 | 0 | 1 | 99.20% | 53.10% | 1.59% | 2.82% | 3.53% | 74.40% | 2.67% | 55.60% | 7.29% |
| SVF#8 | 11 | 23.5 | 92% | 65.50% | none | none | none | 17% | none | none | none |
| SVF#9 | 0 | 1 | 99% | 33.90% | 5.30% | 9.44% | 2.72% | 56.70% | 4.50% | 27.20% | 5.17% |
| SVF#9 | 4 | 6 | 95.50% | 74.50% | none | none | none | none | none | 23.20% | 11.60% |
| SVF#9 | 14 | 20 | 97.30% | 82.60% | none | none | none | 44.70% | none | none | 1.11% |
| SVF#10 | 15 | 20.5 | 96.90% | 89.20% | none | none | none | 5.94% | none | 2.27% | 1.54% |
| SVF#10 | 15 | 24 | 94.50% | 77.90% | none | none | none | 15% | none | none | 20.20% |
| SVF#10 | 23 | 27 | 90.40% | 79% | none | none | none | none | none | none | none |
| SVF#11S | 22 | 29 | 95.30% | 65.40% | none | none | none | 6.18% | none | 15.60% | none |
| SVF#12S | 18 | 26.5 | 96% | 78% | none | none | none | 12.80% | none | none | none |
| SVF#14S | 8 | 12.5 | 96.40% | 87% | none | none | none | 9.53% | none | 1.72% | none |
| SVF #3 | 0 | 3.5 | 99% | 26% | none | none | none | 5.14% | none | 1.62% | 2.75% |
