## Supplementary material for "PTEN regulates adipocyte progenitor growth, differentiation and replicative aging": Table S7

**Table S7.** crRNAs used for CRISPR/Cas9 *PTEN* knockout.

| # | Design ID | crRNA Sequence | PAM | Strand |
|---|---|---|---|---|
| 1 | Hs.Cas9.PTEN.1.AC | TTATCCAAACATTATTGCTA | TGG | + |
| 2 | Hs.Cas9.PTEN.1.AF | TATCCAAACATTATTGCTAT | GGG | + |
| 3 | CD.Cas9.VCTN3398.AH | ATATCTGAGTACTTTAGTTA | AGG | - |
| 4 | CD.Cas9.VCTN3398.BE | TTTCCTGCAGAAAGACTTGA | AGG | + |
