## Supplementary material for "PTEN regulates adipocyte progenitor growth, differentiation and replicative aging": Table S8

**Table S8.** Antibodies used for Western blot (Wb) and immunofluorescence staining (IF).

| Primary antibody | Dilution | Distributor | Cat. no |
|---|---|---|---|
| PTEN ( 138G6) Rabbit map | 1:1000 TBS-T 5%BSA (Wb) | CST | #9559 |
| AKT antibody Rabbit polyclonal Ab | 1:1000 TBS-T 5%BSA (Wb) | CST | #9272 |
| Phospho-AKT (Thr308) (224F9) Rabbit mAb | 1:1000 TBS-T 5%BSA (Wb) | CST | #4056 |
| Phospho-S6 Ribosomal protein (Ser235/236) (D57.2.2E) XP® Rabbit mAb | 1:1000 TBS-T 5%BSA (Wb) | CST | #4858 |
| mAb anti -NAMPT Antibody | 1:1000 TBS-T 5%BSA (Wb) | Novus Biologicals | #NBP1-96585 |
| P21 Waf 1/Cip1 D(S60) | 1:1000 TBS-T 5%BSA (Wb) | CST | #2496 |
| alpha Tublin (11H10) Rabbit mAb | 1:2000 TBS-T 5%BSA (Wb) | CST | #2125 |
| Phospho-FoxO1 (Ser256) Antibody | 1:1000 TBS-T 5%BSA (Wb) | CST | #9461 |
| SREBP-1 polyclonal antibody | 1:1000 TBS-T 5% milk (Wb) | Santa Cruz | sc-8984 |
| ANTI-FLAG Rabbit antibody | 1:1000 TBS-T 5%BSA (Wb) | Sigma | F7425 |
| KI-67 (MIB-1) Mouse mAb | 1:200 IF-buffer (IF) | Dako | P0447 |
| **Secondary antibody** | **Dilution** | **Supplier** | **Cat. no** |
| Polyclonal goat anti-rabbit immunoglobulin/HRP | 1:2000 TBS-T 5% milk (Wb) | Dako | P0447 |
| Polyclonal goat anti-mouse immunoglobulin/HRP | 1:2000 TBS-T 5% milk (Wb) | Dako | P0448 |
| Alexa Fluor 488 goat anti-mouse IgG H+L | 1:1000 IF-buffer (IF) | Invitrogen | A11001 |
