## Supplementary material for "PTEN regulates adipocyte progenitor growth, differentiation and replicative aging": Table S9

**Table S9.** Primers used for RT-qPCR.

| Gene | Forward | Reverse | Probe |
|---|---|---|---|
| *HPRT* | GGCAGTATAATCCAAAGATGGTCAA | GTCTGGCTTATATCCAACACTTCGT | CAAGCTTGCTGGTGAAAAGGACCCC |
| *TBP* | TTGTAAACTTGACCTAAAGACCATTGC | TTCGTGGCTCTCTTATCCTCATG | AACGCCGAATATAATCCCAAGCGGTTTG |
| *PTEN* | TGTAAAGCTGGAAAGGGACGA | GGAATAGTTACTCCCTTTTTGTCTC | |
| *ADIPOQ* | GGCCGTGATGGCAGAGAT | CCTTCAGCCCCGGGTACT | CGATGTCTCCCTTAGGACCAATAAGACCTGG |
| *FABP4* | GCTTTTGTAGGTACCTGGAAACTTG | ACACTGATGATCATGTTAGGTTTGG | CCTGGTGGCAAAGCCCACTCCTCAT |
| *FASN* | GGCAAATTCGACCTTTCTCAGA | GGACCCCGTGGAATGTCA | CACCCGCTCGGCATGGCTATCTT |
| *CDKN1A* (*p21*) | CGAAGTCAGTTCCTTGTGGAG | CATGGGTTCTGACGGACAT | |
| *CDKN2A* (*p16*) | CTTCGGCTGACTGGCTGG | TCATCATGACCTGGATCGGC | |
| *CDKN2B* (*p15*) | AGGCGCGCGATCCAG | GGTGAGAGTGGCAGGGTCT | |
| *HIPK2* | GCTCAAGATGGCAGATTCCG | ACTTGCACATGTGAGGCCATA | |
| *FOXO1* | CCTACGCCGACCTCATCAC | AATTGAATTCTTCCAGCCCGC | |
| *RNF144B* | TGCCTGAAACAGTACATGCAG | ACCAAACAGGCAATCTCAGC | |
| *SREBF-1* | ACCGACATCGAAGGTGAAGT | CAGGGAAGTCACTGTCTTGGT | |
